# Fast parametric curve matching (FPCM) for automatic spike detection

**DOI:** 10.1101/2021.12.31.474621

**Authors:** Daria Kleeva, Gurgen Soghoyan, Ilia Komoltsev, Mikhail Sinkin, Alexei Ossadtchi

## Abstract

Epilepsy is a widely spread neurological disease, whose treatment often requires resection of the pathological cortical tissue. Interictal spike analysis observed in the non-invasively collected EEG or MEG data offers an attractive way to localize epileptogenic cortical structures for surgery planning purposes. Interictal spike detection in lengthy multichannel data is a daunting task that is still often performed manually. This frequently limits such an analysis to a small portion of the data which renders the appropriate risks of missing the potentially epileptogenic region. While a plethora of automatic spike detection techniques have been developed each with its own assumptions and limitations, non of them is ideal and the best results are achieved when the output of several automatic spike detectors are combined. This is especially true in the low signal-to-noise ratio conditions. To this end we propose a novel biomimetic approach for automatic spike detection based on a constrained mixed spline machinery that we dub as fast parametric curve matching (FPCM). Using the peak-wave shape parametrization, the constrained parametric morphological model is constructed and convolved with the observed multichannel data to efficiently determine mixed spline parameters corresponding to each time-point in the dataset. Then the logical predicates that directly map to verbalized text-book like descriptions of the expected interictal event morphology allow us to accomplish the spike detection task. The results of simulations mimicking typical low SNR scenario show the robustness and high ROC AUC values of the FPCM method as compared to the spike detection performed using more conventional approaches such as wavelet decomposition, template matching or simple amplitude thresholding. Applied to the real MEG and EEG data from the human patients and to rat ECoG data, the FPCM technique demonstrates reliable detection of the interictal events and localization of epileptogenic zones concordant with independent conclusions made by the epileptologist. Since the FPCM is computationally light, tolerant to high amplitude artifacts and flexible to accommodate verbalized descriptions of the arbitrary target morphology, it may complement the existing arsenal of means for analysis of noisy interictal datasets.

## 1. Introduction

Epilepsy is one of the most prevalent neurological disorders, as it affects around 50 million people worldwide, according to the World Health Organization (WHO). In additional to clinical manifestations such as seizures this disorder is characterized by brief EEG/MEG patterns with a specific morphology – interictal spikes, – produced by synchronous activity of neuronal groups in epileptogenic brain regions (Staley and Dudek, 2006). Interictal spikes are considered to be a valid marker of epilepsy, which can be detected noninvasively and used for the purposes of irritative zone (IZ) localization. The development of algorithms for automatic spike detection is in a great demand due to the fact that the reliability of the standard process of reviewing EEG/MEG data performed by human experts can be low both for the same expert or different experts (Webber et al., 1993; Hostetler et al., 1992; Halford et al., 2013; Scheuer et al., 2017) is usually time-consuming, tedious and often results into incomplete picture.

The existing approaches of automatic spike detection in interictal EEG/MEG include mimetic approaches, morphological methods, template matching, wavelet-based and ICA-based techniques. To date, these techniques are well developed but primarily applicable for detection of salient spikes in the artifact-free data. In what follows, we review these methods and suggest a novel mimetic technique applicable for detection of low-amplitude interictal events in the noisy data with high-amplitude transient artifacts.

The mimetic approach allows for automatization of the heuristics of visual inspection adopted by the experts, increase in sensitivity and reduce the time spent on the analysis of large EEG/MEG datasets (e. g. the data from ambulatory monitoring). One common step that many mimetic approaches share is waveform decomposition accompanied by subsequent evaluation of the parameters of extracted waves or half waves with opposite directions. These parameters may include the sharpness, the amplitude, the duration, the slope and area under the wave (Gotman and Gloor, 1976; De Oliveira et al., 1983; Faure, 1985; Glover et al., 1986; Gotman and Wang, 1991, 1992; Dingle et al., 1993; Keshri et al., 2011; Liu et al., 2013). Such parameters as amplitude, slope and sharpness are usually defined in relation to the background activity, as spike patterns are supposed to be distinguishable from the background. After the features of waves are extracted, the classification is performed with regards to the predefined thresholds derived from the expert’s knowledge. Certain implementations of mimetic algorithms are accompanied by capturing the state transitions (Faure, 1985; Keshri et al., 2011); by detection of non-epileptic transients depending on preliminarily classified states (e.g. active wakefulness or slow wave sleep) (Gotman and Wang, 1991, 1992) and by the use of spatial context (Dingle et al., 1993) and additional EKG, EOG or EMG data for elimination of false positives (Glover et al., 1989).

Another group – morphological methods – is based on the principles of mathematical morphology and four basic operations: erosion, dilation, the combination of which gives opening and closing operations (Maragos and Schafer, 1987). Applied to a given time series, opening and closing smooth the signal from below and from above correspondingly, allowing for cutting down the spikes and filling up the valleys between them. Nishida et al. (1999) used polynomial functions of the second order as the structuring elements for closing and opening operations in order to detect spike portions and slow portions of the spike waveforms. The minimization of the cost function was used to determine the parameters of the structuring elements. Since spikes can be bi-directional, symmetrical polygons (e. g. a unit circle disk) were proposed as the structuring element (Pon et al., 2002). The lack of analytical justification for the choice of the structuring elements (in particular, their amplitude and width) motivated Xu et al. (2007) to introduce an optimization criterion that is sensitive to the rate of extraction of spikes and suppression of background activity and defined as the ratio of maximal amplitude divided by the mean to the number of zero-pass points divided by the length of the signal.

Taking into account significant between subjects variability, the methods based on template matching became a compromise between time-consuming manual spike detection and fully automatic approaches. These methods operate on the set of manually selected spike templates in order to perform a comparison with the given data segment. The similarity between the template and the data can be evaluated by the means of correlation (Kim and McNames, 2007; Lodder et al., 2013), MSE test (El-Gohary et al., 2008), Euclidean distance (Sankar and Natour, 1992) and other metrics. The templates can be patient-dependent (taken from the recording of a given patient) or patient-independent (built on the basis of the spikes from multiple patients) (Sankar and Natour, 1992). On the one hand, if the database is large enough (Lodder et al., 2013) the latter type of templates is more immune to the variability of spike morphology. On the other hand, patient-dependent templates can increase the sensitivity of the algorithm to the within-patient spike properties (Sankar and Natour, 1992). Nevertheless, even within one patient the algorithm based on the patient-dependent templates (which are usually chosen from the first spikes of the recording) may not be reproducible, since the intrapatient variations cannot be excluded. In this case, instead of using manual construction of templates, Nonclercq et al. (2009) proposed to average automatically detected spikes to obtain a patient-specific template for tailoring of the detection algorithm to the patient. In order to strike a balance between intrapatient variability and patient specificity, the averaging of the preliminarily detected spikes can be replaced by clustering of events with similar morphological properties (Nonclercq et al., 2012).

The morphological uniqueness of spikes can also be maintained through the use of multichannel templates (Ji et al., 2011). A bipolar montage is used based on the polarity, amplitude, and distribution of all bipolar spikes. Finally, spike events, clustered by focal channels, are used to form a multichannel pattern: extraction is performed only for the channels with the sufficient number of events.

Considering the non-stationary nature of spikes, wavelet analysis was suggested to automate spike detection(Senhadji et al., 1995). While single-level wavelet analysis(Sartoretto and Ermani, 1999) is applicable to the data with distinguishable transients and little artifacts, a multi-resolution and multi-level wavelet approaches allow differentiation of spikes from non-epileptoform events (Calvagno et al., 2000; Indiradevi et al., 2008). In general, wavelet decomposition can be followed by detection based on the constant threshold (Sartoretto and Ermani, 1999), adaptive thresholds (Indiradevi et al., 2008), sub-band specific thresholds (Calvagno et al., 2000) or more sophisticated approaches employing artificial neural networks (Subasi, 2006)). Moreover, Latka et al. (2003) studied the behaviour of the wavelet transform as a function of scale and used it as a detection criterion.

The described groups of approaches operate on the timeseries level and are not explicitly aimed at localizing the sources that generate interictal spike activity and do not take into account the information about spatial properties of the cortical generators of interictal spikes. Methods, based on ICA (independent component analysis)-decomposition of multi-channel recordings, are used for identification of independent components characterized by spike activity and subsequent localization using inverse mapping (e. g. RAP-MUSIC (Recursively Applied and Projected Multiple Signal Classification) (Mosher and Leahy, 1999)). While some methods require visual inspection of independent components for extraction of ’epileptic subspace’ (Kobayashi et al., 2002a,b), the others suggest objective measures, such as ’spikyness index’ defined as the ratio of absolute maxima to the mean absolute value of the time series (Ossadtchi et al., 2004). In order to avoid manual exclusion of spurious sources, Ossadtchi et al. (2004) proposed the use of spatio-temporal clustering applied to the localized sources and retaining only statistically significant clusters.

In this paper we describe a novel method that we dub as fast parametric curve matching (FPCM) for automatic spike detection, which is based on the constrained mixed spline model and adopts the principles of mimetic approaches, while being scale-independent. FPCM is simple to implement and has low computational demands. Within this method, the peak-wave shape is parameterized by two linear segments and one second-degree polynomial. The coefficients of this structure are obtained as the result of convolution of this morphological model and the data. The process of decision making is defined by logical predicates on signs and ratios of the coefficients. We report on realistic simulations used to evaluate performance by computing receiver operating characteristic (ROC) curves and areas under them. We also present the application of FPCM to 4 clinical datasets and the data received from the rat. We then discuss the advantages and weak points of the proposed solution and outline the directions for its further development.

## 2. Methods

### 2.1. Description of the approach

We parameterize the shape of an interictal spike-wave complex *s*(*t*) by a three-segment mixed spline with two linear and one second-degree polynomial as follows:

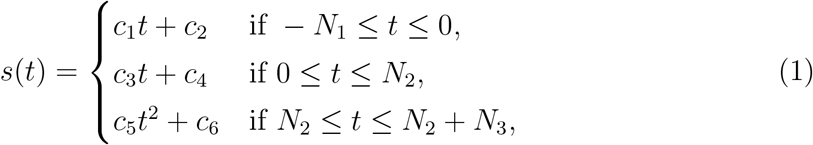

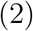

so that *c*_1_ and *c*_3_ are the peak’s left and right slopes, *c*_2_ and *c*_4_ are the corresponding intercepts, *c*_5_ scales the parabolic curve approximating the wave part of the interictal discharge and *c*_6_ is wave’s intercept (see Fig. 1). Without the loss of generality we assume that the peak of the spike corresponds to *t* = 0. We also impose the continuity constraints to guarantee no jumps between the segments in our model, i.e. *s*(0_) = *s*(0_+_) and *s*(*N*_2−_) = *s*(*N*_2+_) where ”-” and ”+” denote the left and the right neighborhoods correspondingly.

**Figure 1:**
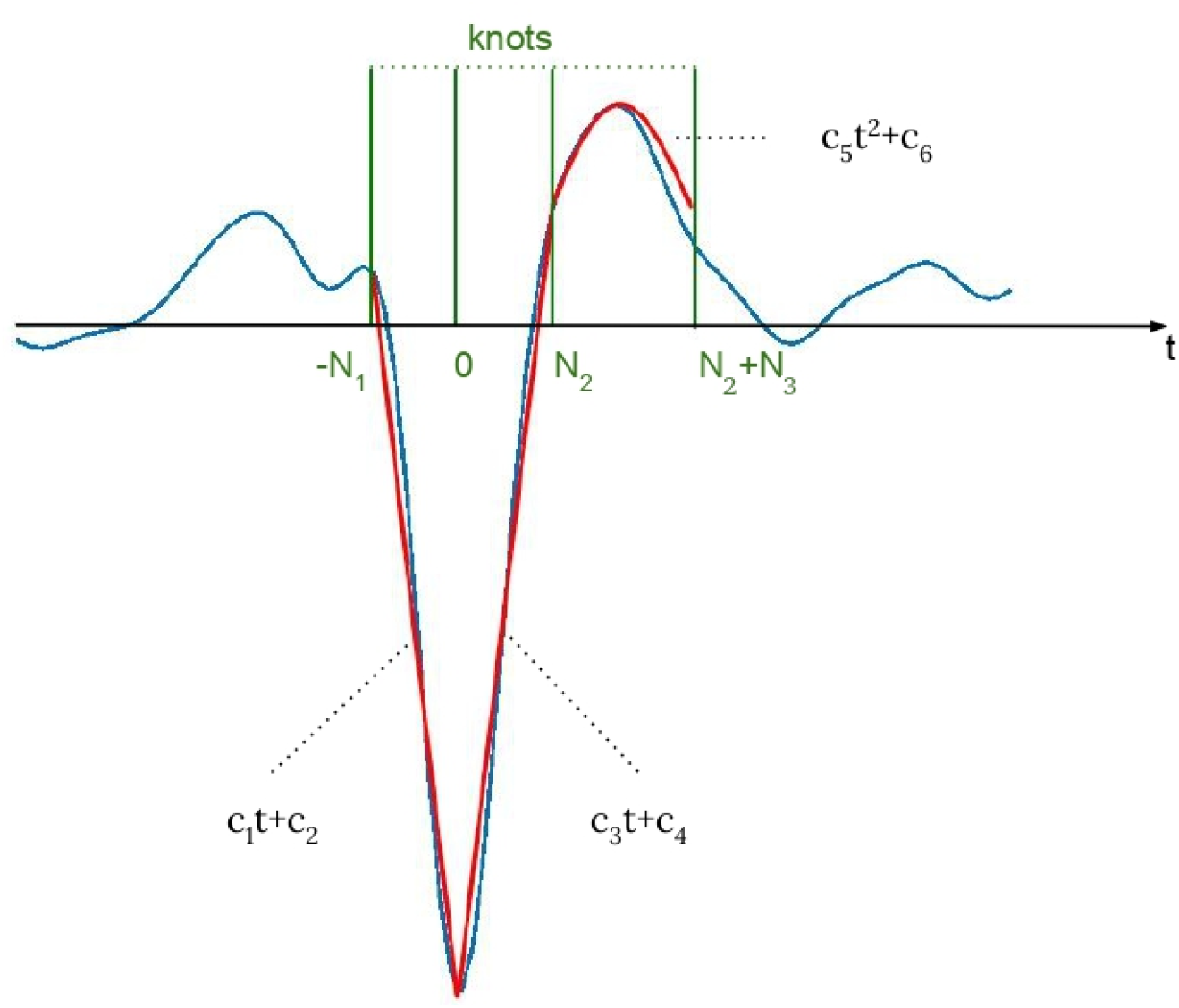
The parametrization of the spike shape (blue) with the spline (red)

This mixed spline model allows us to efficiently capture the morphological properties of the timeseries segment and mimic the process of manual spike data analysis. For that, after fitting spline parameters we employ the corresponding logical predicates that facilitate automatic human-like decision making regarding the timeseries morphology in each N-samples long sliding window, *N* = *N*_1_ + *N*_2_ + *N*_3_ + 1. As we show below the fit of the constrained mixed spline model can be efficiently performed via a few convolution operations between the channel timeseries data and N-sample long rows of the inverted mixed spline model which makes the proposed approach efficient in analysing large amounts of data and feasible for a real-time implementation.

To fit the model (2) we minimize the residual quadratic error. In its simplest form the coefficients of the polynomial spline correspond to the solution of the following equation:

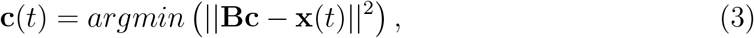

where **x**(*t*) = [*x*(*t*–*N*_1_), *x*(*t*–*N*_1_ + 1),…, *x*(*t*+*N*_2_+*N*_3_)]^*T*^ is a single channel data segment centered around the peak of the spike’s sharp wave, **c**(*t*) = [*c*_1_(*t*),…, *c*_6_(*t*)]^*T*^ is the vector of resulting spline coefficients for data segment **x**(*t*), and **B** is the morphological model matrix.

We construct our mixed spline morphological model matrix **B** as follows. First, we define three elements of the forward model corresponding to the three elementary segments (down-slope, up-slope and wave):

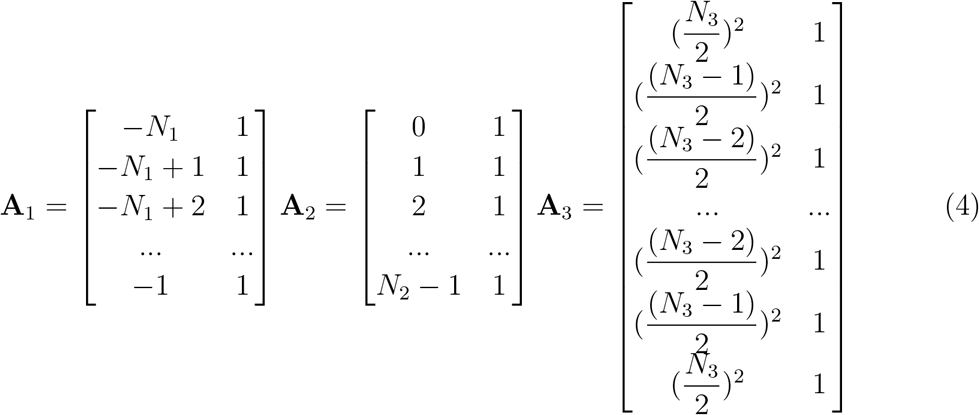

where **A**_1_ and **A**_2_ correspond to the linear segments and **A**_3_ – to the wave. The first column of each matrix represents the abscissa raised to the spline power, where *N*_1_, *N*_2_ are the predefined duration in samples of the down and up slopes of the sharp and *N*_3_ is the duration of the slow wave. The entire spline comprising three segments can be represented as **x** = **Ac** by scaling these abscissa values with coefficients from vector **c**. In case of the mixed spline model considered here our unconstrained morphological model is given simply by the block diagonal matrix **A**:

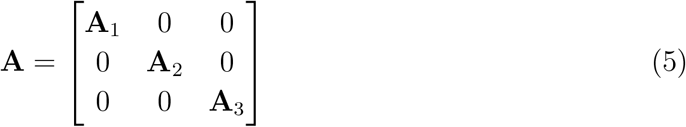

**A**_1_, **A**_2_ and **A**_3_ independently represent three elements of the spike. In order to make the mixed spline coefficients meaningful we must implement the continuity constraints mentioned earlier. The last point of the first linear segment must coincide with the first point of the second linear segment, while the last point of the second linear segment must coincide with the first point of the wave segment. To do so, after putting down the corresponding constraint in the form of two equations we can project the spline matrix away from the 6 dimensional vectors made of the coefficients in the constraint equations. More specifically, requiring that **A**_1_(*end*, 1)*c*_1_ + **A**_1_(*end*, 2)*c*_2_ = **A**_2_(1, 1)*c*_3_ + **A**_2_(1, 2)*c*_4_ for any values of *c*_1_, *c*_2_, *c*_3_, *c*_4_ and analogously enforcing the second constraint we arrive at the constraint vectors:

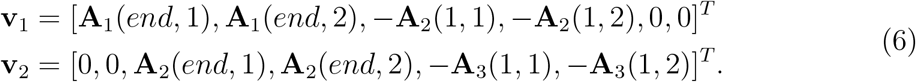

In the above we used Matlab notation to naturally refer to the first and to the last rows of the elementary matrices **A**_1_,**A**_2_ and **A**_3_ of our morphological model.

Satisfying the above mentioned first order continuity constraints is equivalent to ensuring that vectors **v**_1_ and **v**_2_ belong to the right null-space of the morphological model matrix. This can be achieved by projecting the rows of **A** into the subspace orthogonal to **v**_1_ and **v**_2_ which yields the constrained mixed spline model matrix **B**:

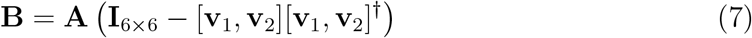

Note that it is possible in a similar fashion to introduce additional constraints and match, for example, the derivatives over both positive and negative neighbourhoods of the peak-wave junction, corresponding to *t* = *N*_2_, see Fig. 1.

Then, solving (3) for the vector of spline parameters **c** in the least squares sense via Moore-Penrose pseudo-inverse of **B** we get the following expression for the mixed spline coefficients:

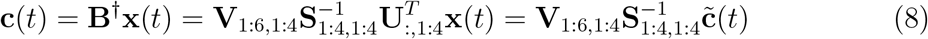

where matrices **U**,**S** and **V** are such that **B** = **USV**^*T*^ and obtained as the SVD (singular value decomposition) of **B**. The subscripts denote the used range of rows and columns of these matrices. Vector 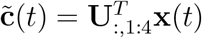 is a vector of latent coefficients.

For a long input sequence *x*(*t*), the elements 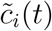 of 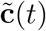 for each time instance *t* can be computed by convolving *x*(*t*) with the first four filters formed from the first four rows of **U** whose elements are reversed along the vertical (time) dimension. We will denote these flipped vectors as 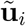, *i*,..,1,4 and then

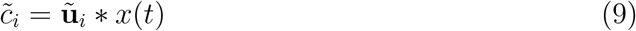

where * denotes convolution operation.

Then, the obtained vector of latent coefficients can be transformed to the original and interpretable spline coefficients vector **c**(*t*) according to (8). The use of SVD on the one hand is instrumental in computing Moore-Penrose pseudo-inverse of rank-deficient matrix **B** and on the other allows us to reduce computations when applied to the actual data in a sliding window mode. According to (8) in order to compute the vector of six spline coefficients we can perform only four convolutions due to the linear dependency in the rows of **B** introduced by the projection (7) and then transform the vector of latent coefficients 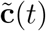 into the six dimensional space of the original spline coefficients **c**(*t*) according to (8).

The procedure described above allows us to very efficiently obtain values of spline coefficients **c**(*t*) for every time point in the original data. Whether or not a data segment [*x*(*t* – *N*_1_),…*x*(*t*),…*x*(*t* + *N*_2_ + *N*_3_)] around some specific *t* exhibits the spike-wave morphology can be established using a set of logical predicates on the elements of vector **c**(*t*) as they carry information about all morphological properties of each segment given that the residual error is low.

The residual error timeseries vector for each segment centered around some time moment *t* can be easily computed as **e**(*t*) = **x**(*t*) – **Bc**(*t*) for all time points at once. Based on our experiments we found that it is beneficial to separately assess the residual scalar error corresponding to the peak and wave segments of the spike. Therefore we introduce two error terms

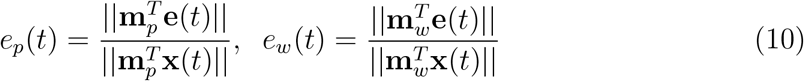

where 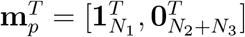 and 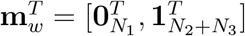 are the corresponding peak and wave binary mask vectors. Note that scalar data vector norms 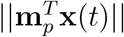 and 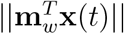 representing the denominator of the relative residual error values *e_p_*(*t*) and *e_w_*(*t*) can also be efficiently computed for all values of *t* using convolution of *x*(*t*) and the two box-car filters of length *N*_1_ and *N*_2_ + *N*_3_ correspondingly.

Observing sufficiently low residual error at time sample *t*, i.e. *e_p_*(*t*) < *θ_p_* and *e_w_*(*t*) < *θ_w_*, indicates that the constrained spline model appears to represent well the timeseries morphology over [*t* – *N*_1_,*t* + *N*_2_ + *N*_3_] interval. The decision making about the presence of an interictal spike at sample *t* is then based on the logical predicates on signs and ratios of the elements of the spline coefficients vector **c**(*t*). Also, using these coefficients we can effortlessly estimate the spline values at some key points and employ them within the logical predicate. For example, as described next we found that peak and wave relative amplitude values are the crucial parameters defining the characteristic shape of the pathological interictal peak-wave complex. These values can be directly inferred from the coefficients for each time point as *h_p_*(*t*) = *c*_4_(*t*) and (*t*) = *c*_6_(*t*) correspondingly.

Each individual user of the proposed approach can employ a specific set of logical predicates based on his or her own experience. We will next describe as an example a list of possible predicates that we used in this study. The predicates are created naturally based on the verbal description of the expected spike morphology as outlined in Table 1.

**Table 1:**
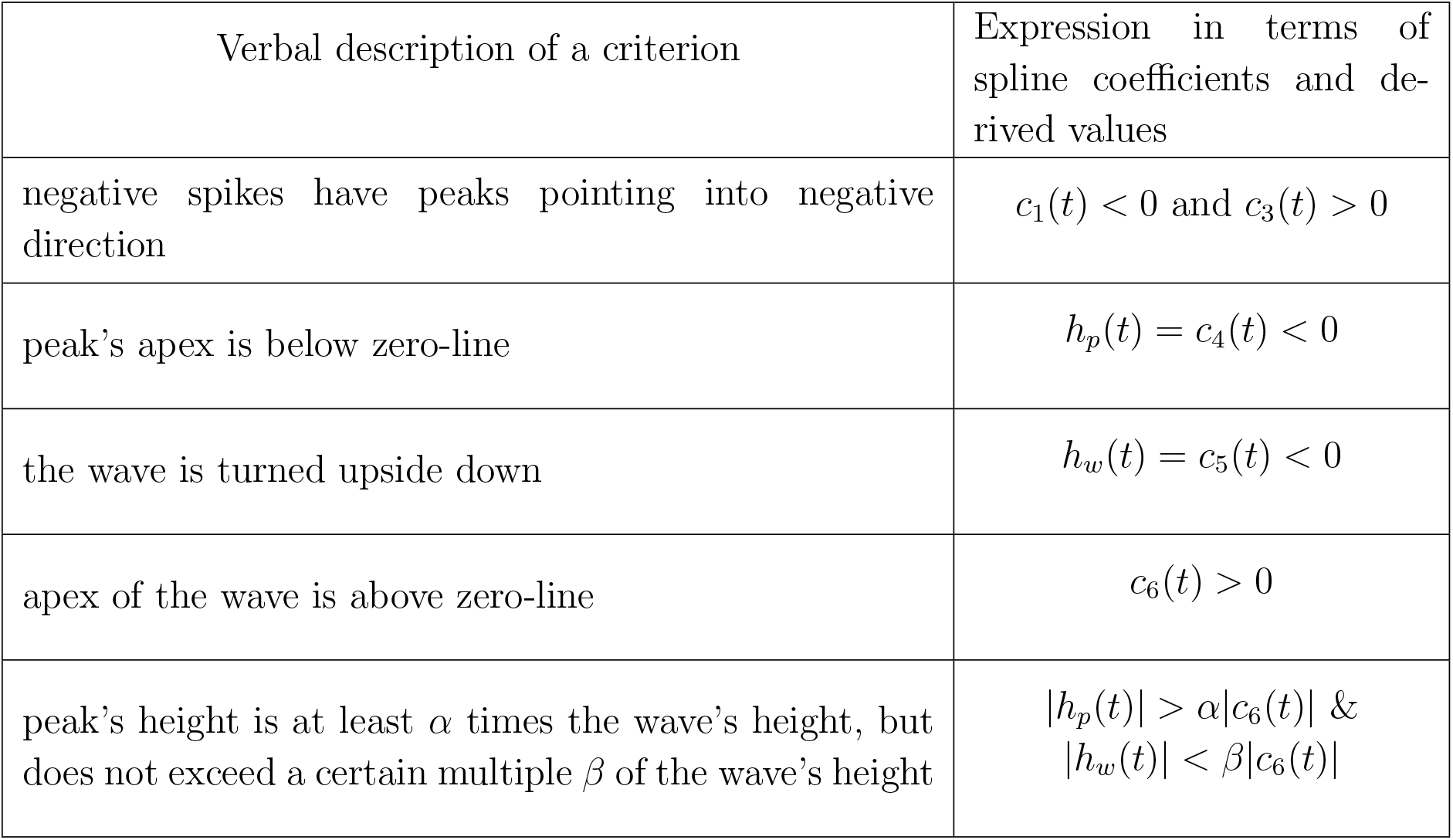
Example of logical predicates on spline parameters (right column) corresponding to the verbal statements regarding spike’s morphology (left column) for a spike pointing into negative direction.

Based on the corresponding clinical literature highlighting e.g. Komoltsev et al. (2020); Kural et al. (2020) spike salience is another important parameter used by human experts when detecting epileptogenic activity. The extent to which a spike stands out of the local background can be easily computed, for example, as the ratio of the absolute peak value, *h_p_*(*t*) = *c*_4_(*t*), to the standard deviation of the background activity within the time window preceding and succeeding the considered spike time interval. The standard deviation values can be efficiently computed by convolving the squared timeseries with a boxcar sequence normalized by its length.

Finally, in the manual analysis of interictal there is an additional requirement that the spike is present on several channels (Adjouadi et al., 2004). That is why, despite the fact that FPCM is consequently applied to single-channel data, we implement this by requiring the criteria described above to be satisfied in at least *N_min_* channels for a single time point *t* or within its immediate *δ* neighbourhood. Once this is the case, we decide that an interictal spike is present at *t*. *N_min_* can be adjusted depending on the criteria provided by the clinician for each particular case. Typically we used *N_min_* = 4 for the MEG data with 102 or 204 channels.

The pipeline described above is briefly presented in Fig. 2.

**Figure 2:**
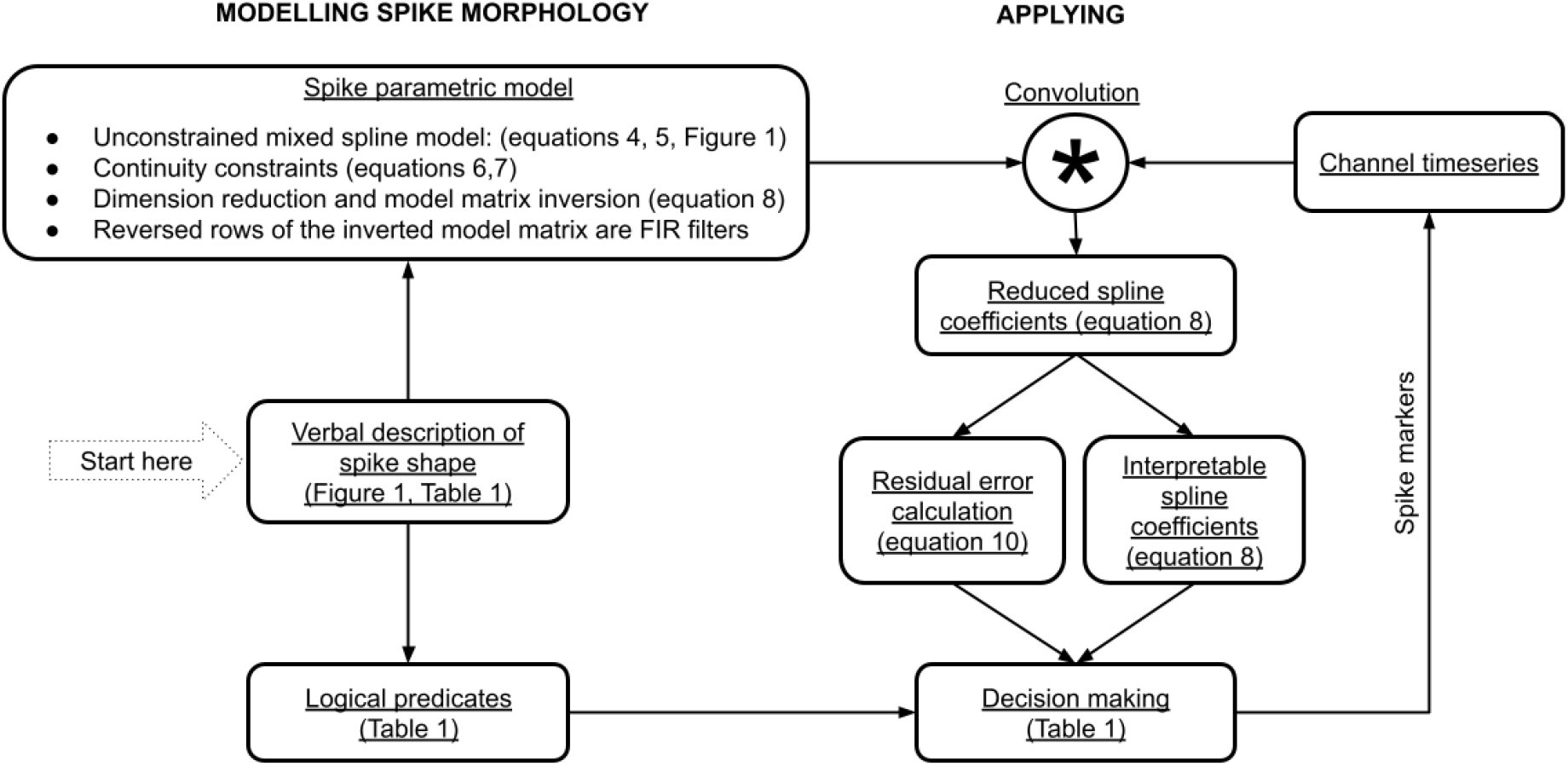
The general steps of implementing FPCM. Based on the verbal description a mixed spline model is created and a set of logical predicates is formulated. After projecting the mixed spline model matrix away from continuity constraints equations introducing the continuity constraints, the spline model matrix is inverted using SVD and dimension reduced FIR filters are formed. These filters are then applied to timeseries via standard highly computationally efficient convolution operation. This yields reduced model spline coefficients for each time slice. After that the reduced spline coefficients are converted back to the vector of interpretable coefficients **c**(*t*) and the residual error along with logical predicates is evaluated for each time slice. Both operations are easily vectorized for efficiency.

### 2.2. Temporal scale invariance

To find the optimal values of *N*_1_, *N*_2_ and *N*_3_ we suggest to use a simple grid-search over a set of the values and their ratios. Then, we simply assess the count of detected spikes and chose the corresponding combination of sub-segment duration values to obtain the final set of spike markers.

An illustration of this procedure is given in Fig. 3 where we show the total count of detected events with the expected morphology as a function of the scale parameter. Given that the procedure of computing spline coefficients **c** is fast, the described scan over temporal scales can be accomplished seamlessly. While in principle for each value of the temporal scale parameter it is possible to perform a full blown analysis including source localization and source-space based clustering, in our work we have used the scale corresponding to the maximum of spikes count curve as shown in Fig. 3. All results on the performance characteristics were obtained in this scale-invariant fashion using the best scale parameter automatically inferred from the actual data using the spikes count curve.

**Figure 3:**
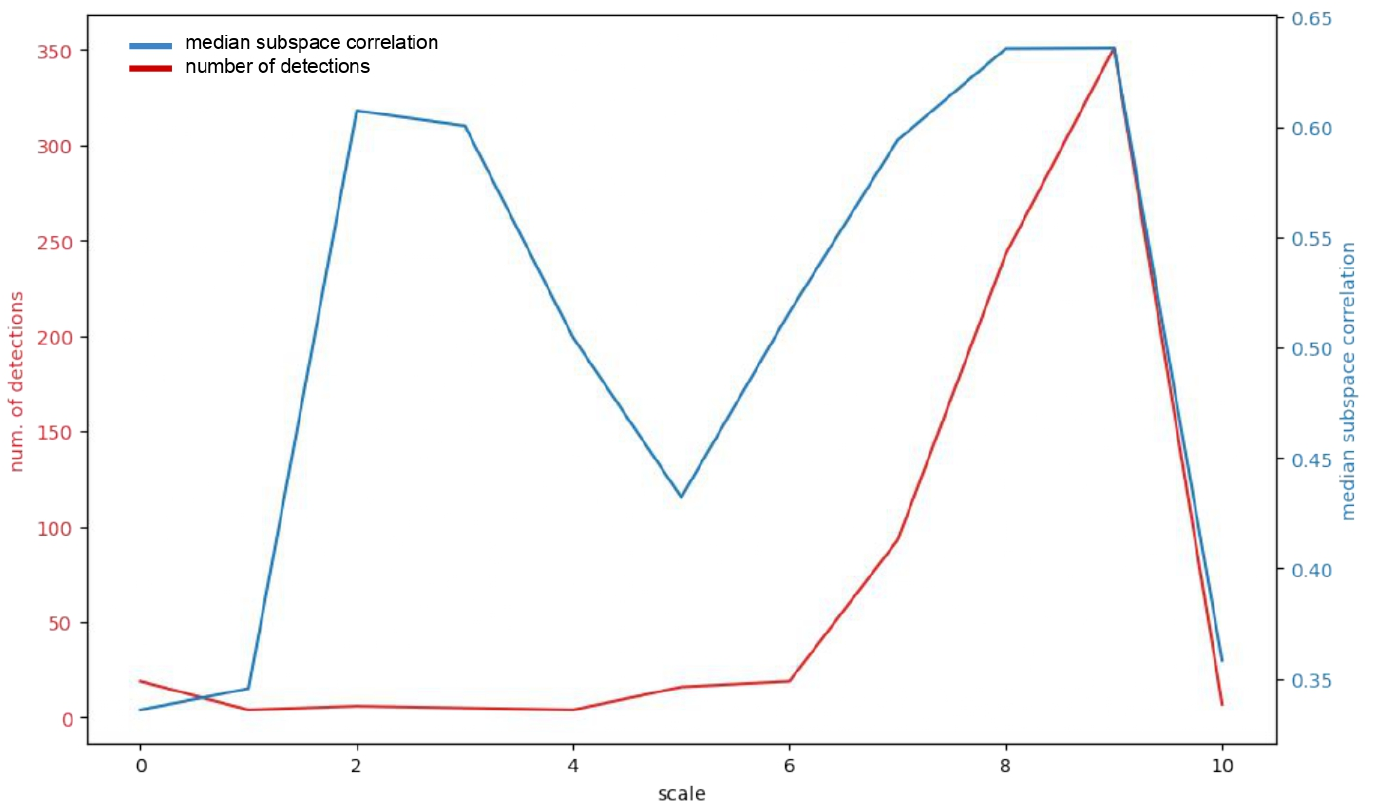
The total count of detected events as a function of the scale parameter and the corresponding median subspace correlations (Mosher and Leahy, 1999) of the fitted dipoles. The first peak in the median subspace correlation curve corresponds to detecting a small portion of true spikes. Scale increase implies wider wave shapes, less true spikes are detected resulting in the decrease of dipole fitting performance. As the scale, at which the maximal number of true spikes is detected, is approached, the subspace correlation predictably increases. The peak in the number of detected events corresponds to *N*_1_ = 10 ms, *N*_2_ = 10 ms and *N*_3_ = 95 ms.

The development of effective methods of scanning through *N*_1_, *N*_2_ and *N*_3_ can supply FPCM with additional flexibility and scale-invariance.

### 2.3. Spike localization

Localization of potentially epileptogenic zones is the main goal of interictal spikes analysis. Therefore, our next step is the use of RAP-MUSIC algorithm (Mosher and Leahy, 1999) for localization of equivalent current dipoles from the spike data in the range from −20 ms to 30 ms around the peak of the detected spikes.

When dealing with MEG data, in order to find source location **r**\* RAP-MUSIC matches signal subspace spanned by the doublet of topographies **G**(**r**) = [**g**_*x*_(**r**), **g**_*y*_ (**r**)] of two orthogonal sources at **r** and the signal subspace extracted from the data matrix **X** around each spike that we estimate as the subspace spanned by the first *R* left singular vectors **U**_*R*_ = [**u**_1_,…, **u**_*R*_] of **X**.

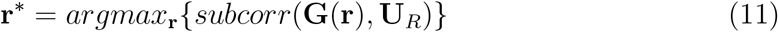

Then, after finding the optimal orientation vector *θ* of the dipole at **r**\* we project both the data and the forward model away from the oriented topography **g^*θ*^** (**r**\*) = **G**(**r**\*)*θ* of the optimally oriented dipole at **r**\* and repeat the scan. We stop when the achieved subspace correlation values falls below the user specified threshold. Subspace correlation reflects the similarity between the subspaces spanned by the columns of the two matrices. If subspace correlation is equal to 1, then the two subspaces share at least one subspace. On the other hand, if subspace correlation is equal to 0, the subspaces are orthogonal. Using this recursive operation RAP-MUSIC resolves the potential problem of the proximity of local maxima in the subspace correlation profile. Typically, when working with real data we find one or two dipoles per interictal spike.

### 2.4. Clustering

After having obtained a set of equivalent current dipole locations corresponding to the detected interictal spikes we perform clustering by means of a simple multi-start K-means procedure to identify a set of cortical irritative zones.

### 2.5. Simulations

In order to validate the proposed method we performed a set of realistic simulations. For the sake of estimating the robustness of the algorithm with respect to the natural variation of the interictal spike shapes, we used three various spike morphologies, shown in Fig. 4 that differ in the relative duration of spike and wave segments. These spikes were taken from the data of a single patient marked by the epileptologist.

**Figure 4:**
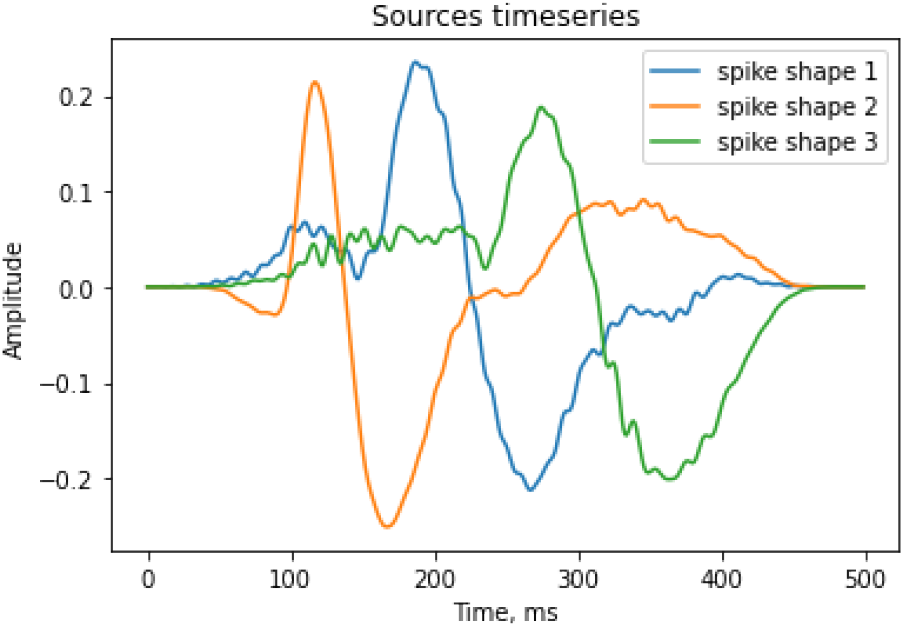
The sample waveforms of the interictal spikes used for realistic simulations.

In the simulated data within each 1000 ms an interictal spike is generated as emanating from one of the three sources located in the right anterior transverse temporal gyrus, left parietal cortex and a deeper source that we placed in the left parahippocampal gyrus. Each spike’s morphology was randomly sampled from the three shapes presented in Fig. 4 and used as a source activation waveform for the equivalent current dipoles that modelled spike generators. The amplitude of each spike was set to result into dipole moment of 70 nAm. We superimposed the simulated interictal activity onto the real sample data from MNE software (Gramfort et al., 2014). The sampling rate of the simulated data was set to 1000 Hz.

Since the real data often contains sharp non-epileptiform transients, which can be erroneously marked as spikes, in order to check the immunity of FPCM to the presence of such artifacts, we supplemented the modeled data with four types of artifacts (see Fig. 5), which occurred 500 ms after the spikes in 80 % of the data (20 % of epochs were left without artifacts). In order to avoid the overlap in further results of dipole fitting, he artifacts were generated by the other set of spatially remote sources: left occipital cortex and orbitofrontal cortex in both hemispheres. By default, the dipole moment of artifacts was set to be equal to the dipole moment of the spikes (70 nAm).

**Figure 5:**
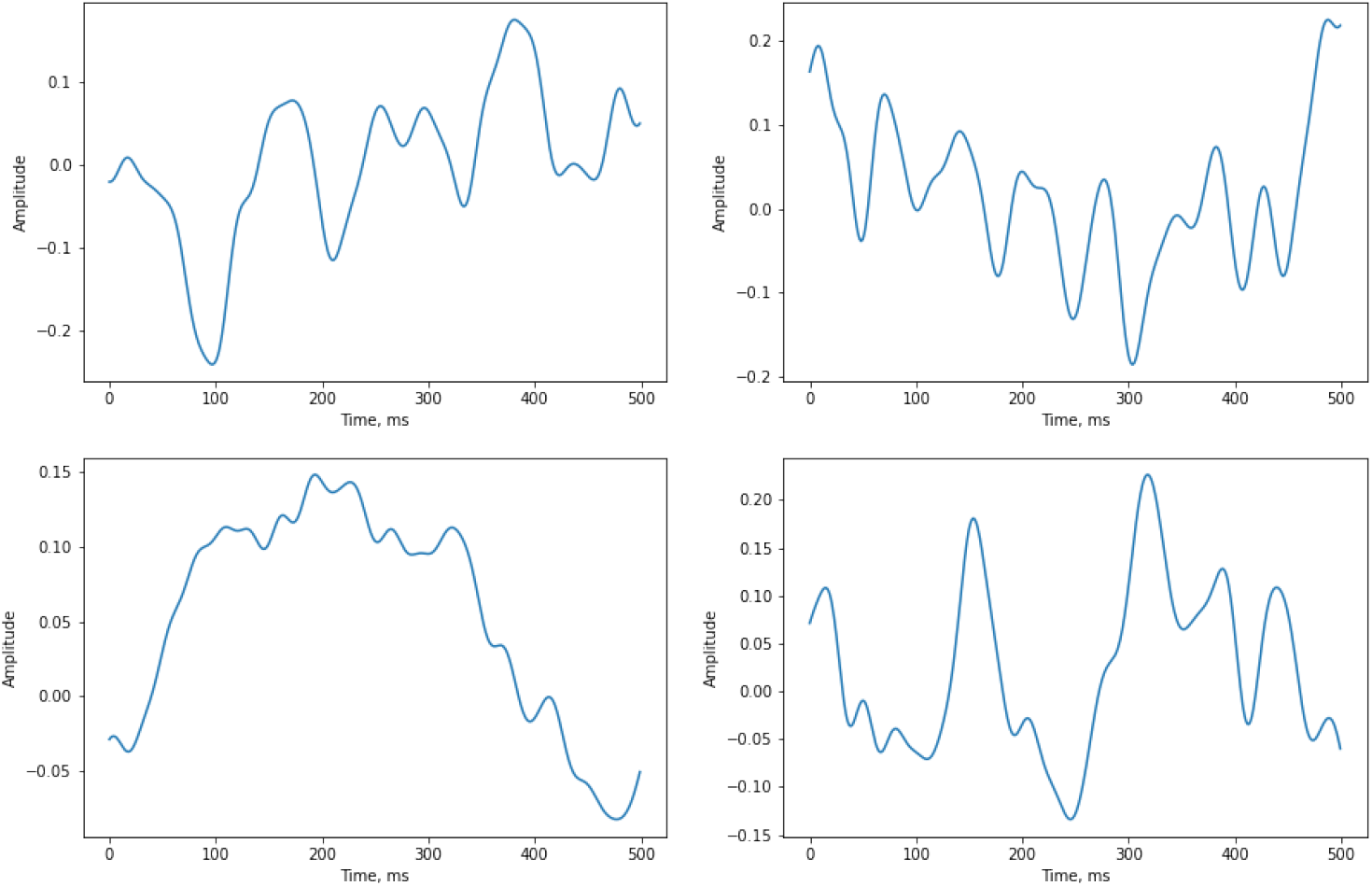
The sample artifacts used for realistic simulations

Additionally, by changing the amplitude of the artifacts from 0 nAm (no artifacts in the whole dataset) to 160 nAm (very expressed artifacts) (see Fig. 6) with the step equal to 40 nAm, we estimated the sensitivity of FPCM’s performance to different signal-to-noise ratios. Thus, SNR was defined as the ratio of the spike’s dipole moment to the artifact’s dipole moment. In this case, we excluded the criterion of the spike’s salience from the set of logical predicates, since it would create a bias in terms of the algorithm’s sensitivity in the case when the artifact’s amplitude exceeds the spike’s amplitude too much.

**Figure 6:**
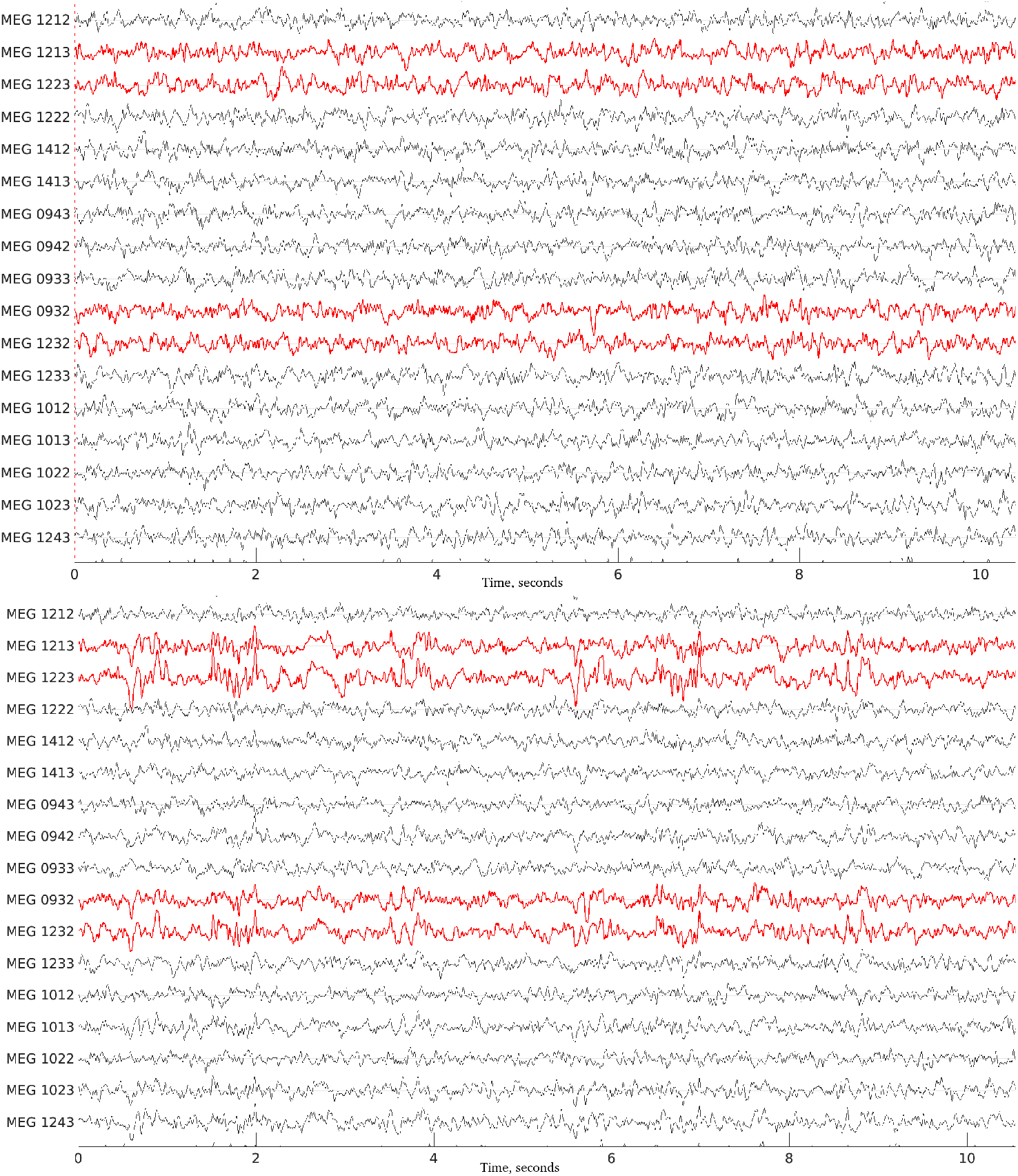
An example of simulated raw data (10-seconds part of it) with the minimal and maximal amplitude of modeled artifacts. The red time series indicate the channels with expressed artifacts

An example of the simulated data is provided in Fig. 7. For each batch in total we generated 600 spikes with approximately 200 spikes per morphology.

**Figure 7:**
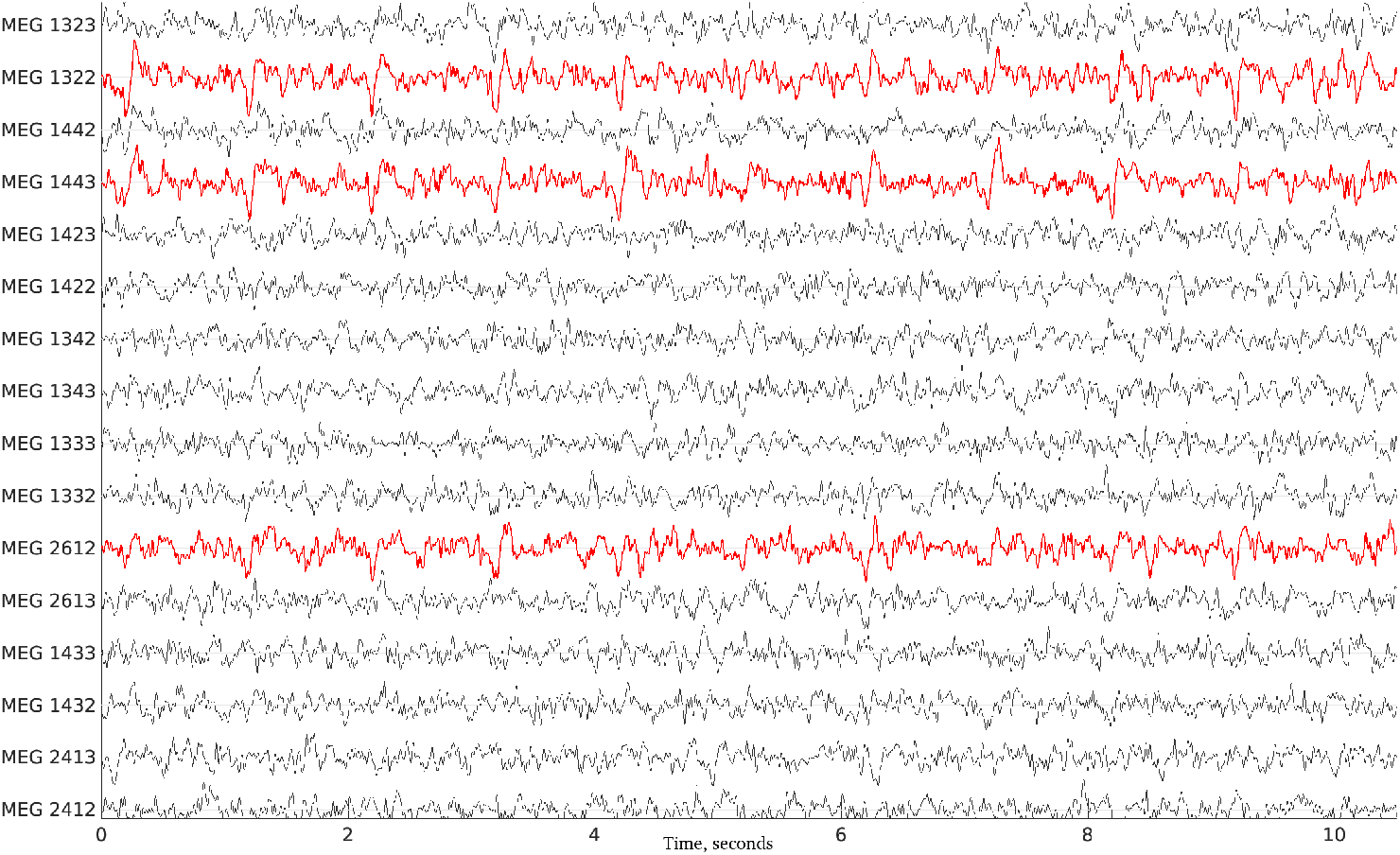
An example of simulated raw data (10-seconds part of it)). The red time series indicate the channels where spikes are present

In the subsequent figures, spike locations were reported only for the events with greater than 65% subspace correlation.

Additionally, we applied the compared methods to the open simulated ECoG dataset, provided by (Quiroga et al., 2004) within testing spike detection and sorting. This data contains single-channel time series with controlled levels of the noise determined from the standard deviation of the signal (from 0.05 to 0.4). The modeled spikes were sampled from the database of approximately 600 average spike shapes from recordings in neocortex and basal ganglia. The morphology of those spikes formed three distinct groups, as the spikes had different scales. The sampling rate of the data was equal to 24000 Hz.

### 2.6. Real data

In order to assess the validity of the proposed approach, we have used 3 MEG datasets from 3 different subjects with diagnosed epilepsy. The first two datasets were recorded using a whole cortex CTF neuromagnetometer with 68 sensors (1st order co-axial gradiometers) and 32 reference channels. The sampling rate was 250 Hz. The third dataset was obtained using Neuromag Vector View with 204 planar gradiometers and 102 magnetometers. The original sampling rate was 1000 Hz, but the recordings were subsequently downsampled to 256 Hz.

Before spike detection in real data we performed a set of standard preprocessing steps. The ICA (independent component analysis) was performed and, based on the typical spectral and spatial properties, the artifactual components related to the heart beats, ocular or motor activity were eliminated. Then the data was filtered in the 3 to 40 Hz range. Note that ICA here was used not to select the components containing spikes but rather the components harboring the artifacts commonly present in the MEG and EEG data.

To demonstrate robustness of the FPCM and its ability to detect tiny interictal events in the presence of high amplitude sharp transients we used a 1 minute long segment from a 12-channel clinical EEG dataset recorded from a patient with severe TBI that resulted in epilepsy.

We have also applied the proposed technique to rat’s hippocampus ECoG data following the TBI (traumatic brain injury). The 200s long segment of 6-channel data was manually labeled by an expert with 39 markers corresponding to the interictal discharges.

### 2.7. Performance evaluation

We evaluated performance of our approach using the area under ROC curve (ROC AUC) metrics. To compute the curved we first determine the counts of true positives (TP), false positives (FP), true negatives (TN) and false negatives (FN). The number of false positives corresponded to the number of detected events located further than 100 ms from the nearest true spike. By varying the thresholds of *n* (the first error term *e_p_*(*t*) corresponding to the spike’s peak) we obtained the values of TP, FP, TN and FN to build the parametric ROC curves. The true positive rate (TPR, or recall) represents the proportion of true spikes among all the events that could be possibly detected. The false positive rate (FPR) is the fraction of erroneously detected events with respect to all possible detections. Additionally, we computed precision – the number of TP over the number of TP and FP.

To assess the the reproducibility of our results we performed our analysis on the set of additional ten simulations with the same source dynamics but independent noise samples. We then compute ROC AUC and also report the confidence intervals equal to ± SD, calculated from the bootstrap-based data.

As an additional performance indicator we also present the dependence of TPR on the number of false positives per minute expressed in the number of seconds occupied by the erroneously detected events.

We also compare the performance of the proposed method to the two more standard approaches. The first of them is a wavelet-based approach. In general, this technique implies sub-band decomposition allowing for disentangling the detailed components with sharp signal transitions from the background (for reviews, see (Abd El-Samie et al., 2018; Wilson and Emerson, 2002)). The two parameters of choice that form the basis of this approach is the mother wavelet function and the sub-bands of interest. It was previously reported that a basis wavelet function of Daubechies 4 (DB4) from MATLAB wavelet toolbox matches the morphological properties of spikes with the highest values of correlation (Indiradevi et al., 2008). We have partly replicated the original preprocessing by (Indiradevi et al., 2008), downsampled simulated signals to 256 Hz and decomposed them into six sub-bands: 64-128 Hz, 32-64 Hz, 16-32 Hz, 8-16 Hz, 4-8 Hz, 2-4 Hz, 0-2 Hz. For further analysis we use the fourth sub-band corresponding to the range 8-16 Hz. Then we marked the as a detection the time slices for which the squares of reconstructed wavelet coefficients exceeded a pre-defined static threshold (see Fig. 8) which we varied to sample the ROC curve.

**Figure 8:**
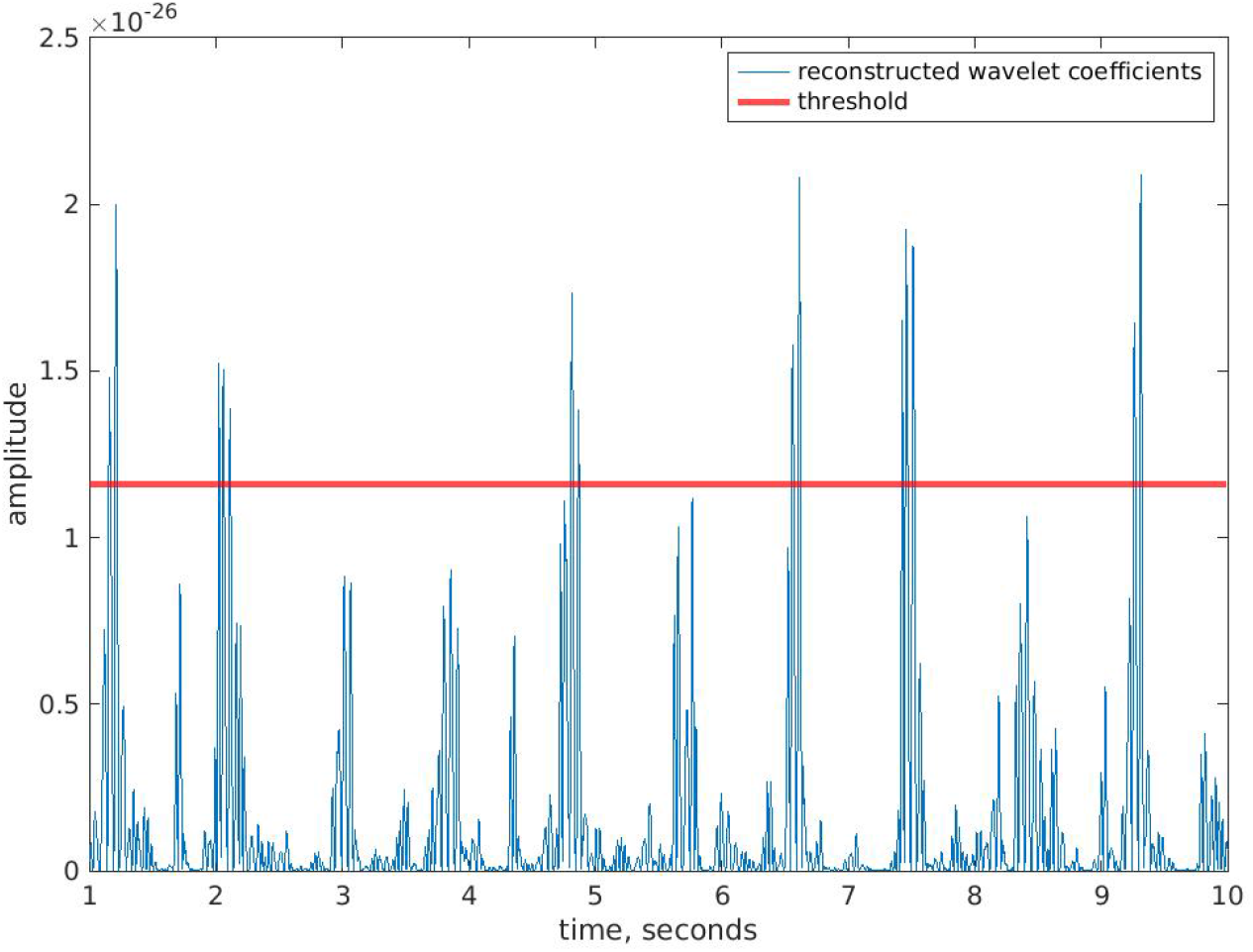
An example of the implementation of the wavelet decomposition and subsequent thresholding in order to detect spike candidates

The second reference method we use for comparison is template matching. We have obtained the templates by manual detections, as in natural circumstances. Then, for each template, the normalized correlation (see Fig. 9) is computed as:

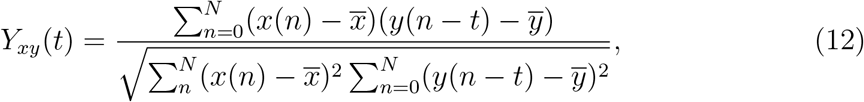

where *x* is the given template, *y* is a one-channel signal, 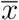 and 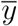 are the averages of the signals over *N* samples, and *t* – is the time corresponding to the placement of the template.

**Figure 9:**
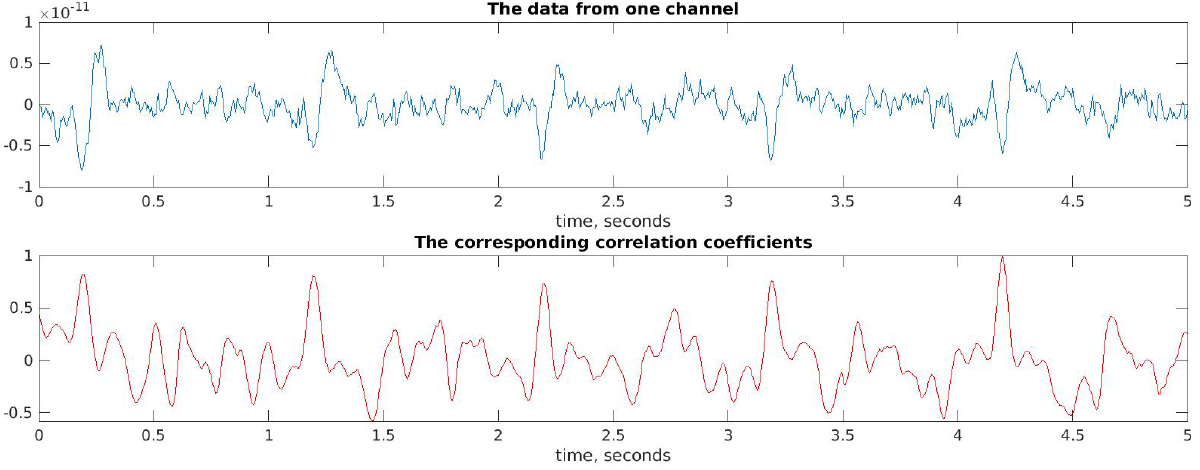
An example of the implementation of the template matching

Those samples which corresponded to the correlation coefficient exceeding the pre-defined threshold were marked as spike candidates.

The programming tools used for implementation of the algorithm and data processing included MATLAB and Python. Specifically, MNE Python Toolbox was used for the realistic simulations and inverse modeling (Gramfort et al., 2014).

### 2.8. Computational complexity

At the core of FPCM there are 6 convolution operations between the short (peak-wave complex length) rows of the mixed-spline model matrix and the channel data timeseries. Therefore the corresponding complexity can be assessed as *O*(*KMN*) where *K* is the number of channels, *M* is the length of the FPCM kernel and N is the timeseries length. Since both wavelets and template matching also rely on convolution, we expect that the FPCM’s computational demands are comparable to that of the other two reference methods. The logical predicate checks are also very well vectorized and therefore can be performed with minimal latency.

## 3. Results

### 3.1. Simulations

#### 3.1.1. MEG

We compared the changes in performance of the proposed method and the standard approaches at different SNR levels by varying the dipole moment of the modeled artifacts: the dipole moment of the spikes was equal to 70 nAm and the dipole moment of artifacts increased from 0 to 140 nAm.

The resulting values show (see Fig. 10) that at low SNRs, when the dipole moment of the artifacts (Fig. 7) does not exceed the dipole moment of the spike generator wavelet based detection outperforms other methods. However, starting from SNR¡=1, when both artifacts’ and spikes’ amplitudes equal to 70 nAm, shape-specific and scale free FPCM outperforms other methods and in general maintains nearly the same performance as with the high SNR data. The superior performance of wavelet decomposition at high SNRs can be explained by the properties of the data itself, since even mere amplitude thresholding without any further processing (see cyan line in Fig. 10) demonstrates even better scores comparable to those achieved with template matching or wavelet decomposition. Another important observation is that the performance of template matching is characterized by high variance across all data realizations. Since the templates were used from the same channels and the same time points per each simulated recording, due to the natural variability of those recordings, the templates differed as well, influencing the performance.

**Figure 10:**
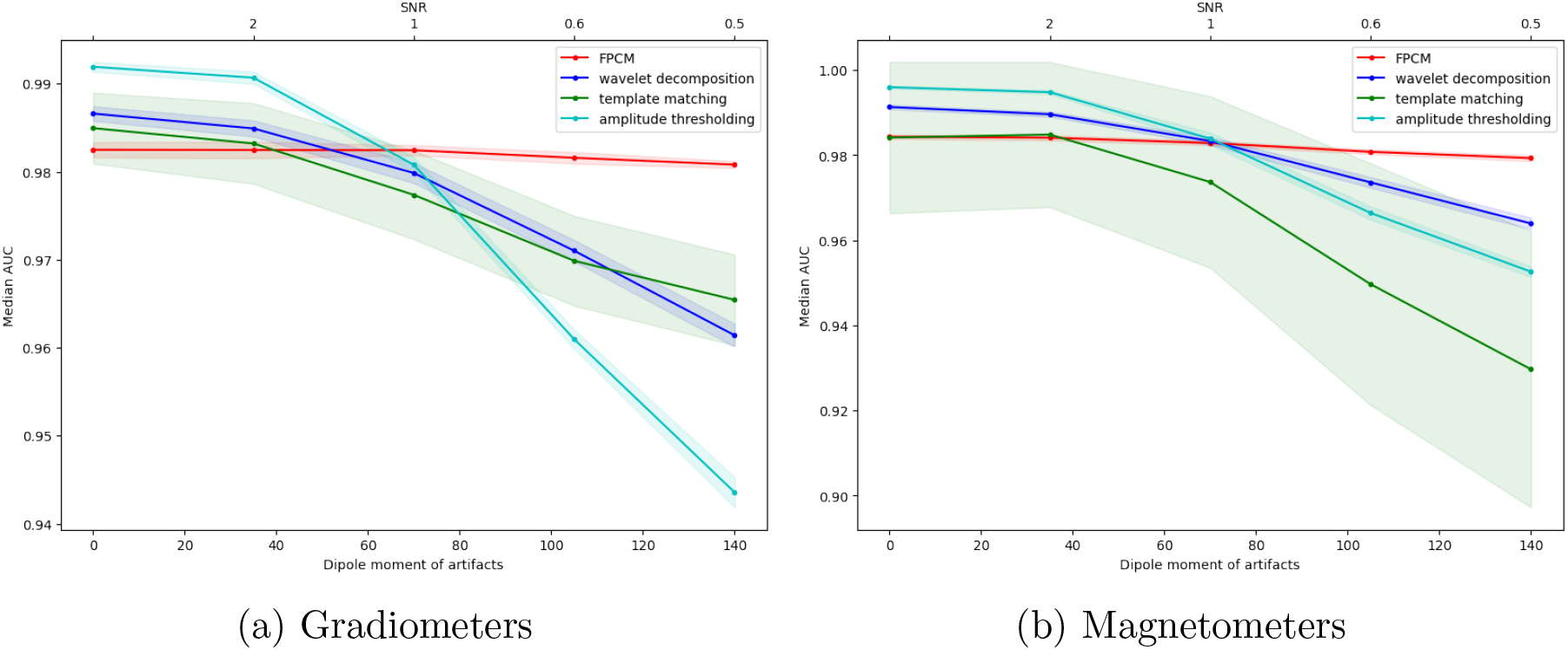
ROC AUC values depending on the SNR. The transparent areas above and below the median AUCs represent the confidence intervals equal to ±*SD*

We have also applied the proposed method of spike detection to the data from realistic simulations with SNR = 1 where we obtained bootstrap-based estimates of the comparative accuracy of the selected methods. Applied to data from gradiometers, FPCM outperformed the standard approaches, demonstrating the highest value of AUC (98.2441) against AUC = 97.9853 for wavelet decomposition and AUC = 97.7368 for template matching (see Table 2 and Fig. 11 (A)). In case of magnetometers, the performance of the compared methods was almost the same: AUC = 98.2871 for FPCM, AUC = 98.3338 for wavelet decomposition and AUC = 97.3692 for template matching. The detailed comparison of AUC values at other SNRs, demonstrating that with its decrease FPCM preserves its high performance, while other methods show the decline in performance, is presented in Appendix in Table 5.

**Table 2:**
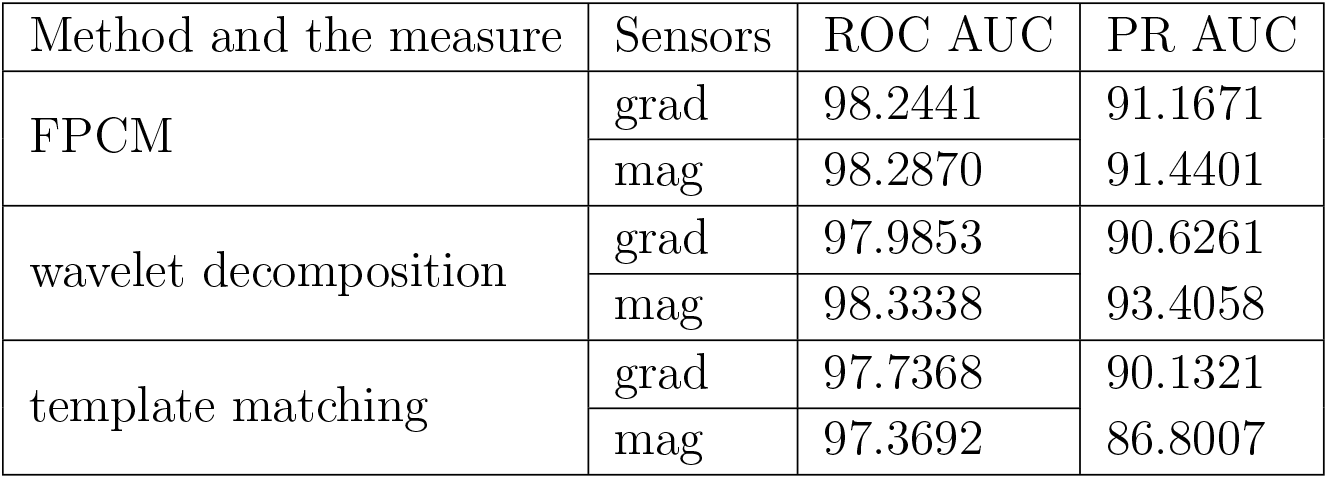
Comparison of ROC AUC values between the methods tested on simulations

**Figure 11:**
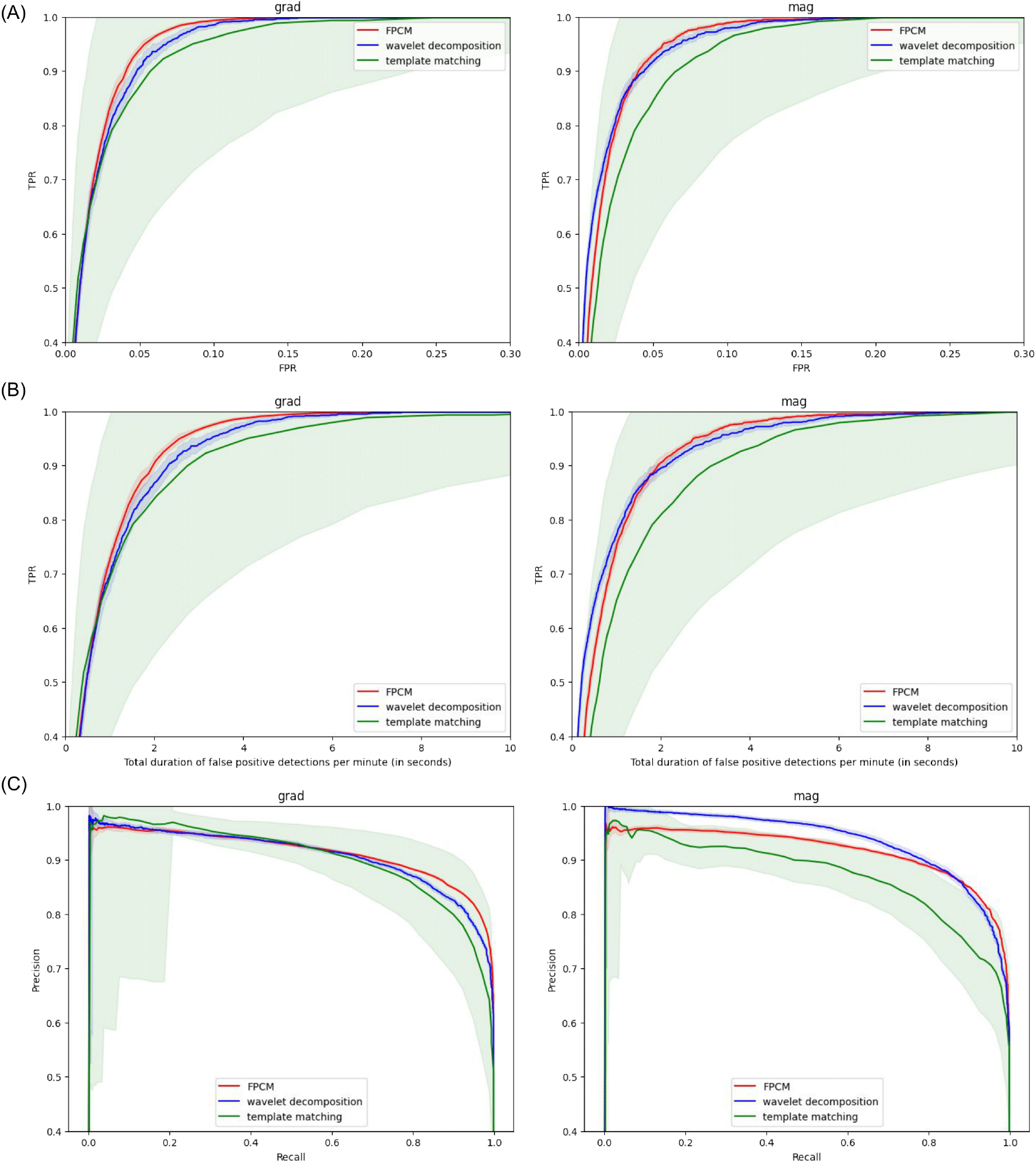
The performance measures of compared methods: (A) ROC curves, (B) TPR vs. total duration of false positives per minute, and (C) precision-recall curves. The transparent areas represent the confidence intervals equal to ±*SD*

The data from ROC AUC is consistent with the additional analysis of false positives per minute, which demonstrates the same tendencies (see Fig. 11 (B)). Finally, precision-recall curves in Fig. 11 (C) show that at high levels of recall FPCM demonstrates superior levels of precision.

Based on this analysis, we found the specific values of thresholds in such a way that it maximizes the difference between TPR and FPR (see Table 3) and assessed the spatial properties of localized spikes for the minimal SNR. Note that in case of the FPCM, the threshold is applied to the spike’s peak error term *e_p_*(*t*), see equation 10.

**Table 3:**
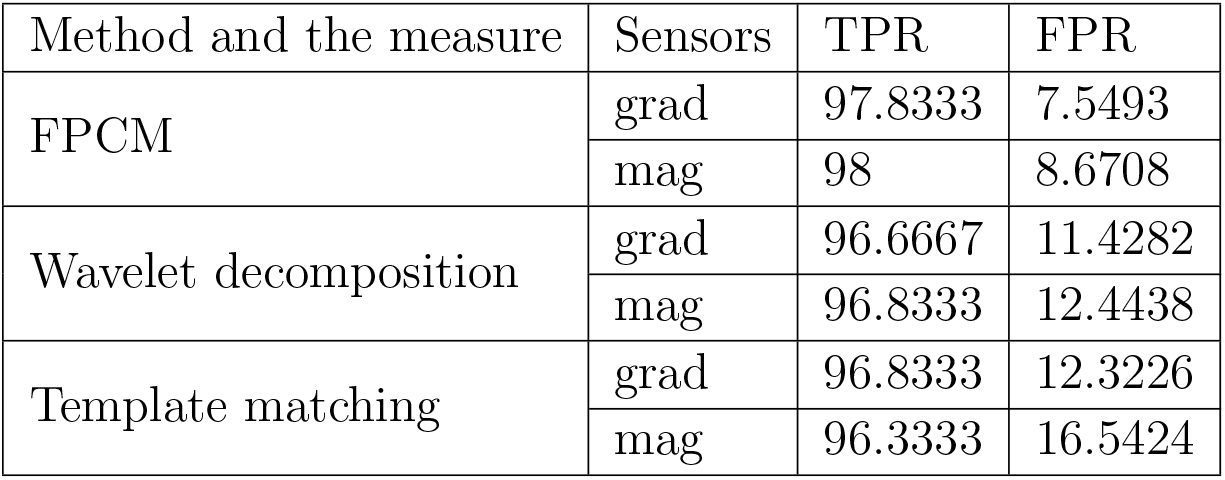
Combination of the TPR and FPR values corresponding to the thresholds used for selection of spikes for source localization.

| Method and the measure | Sensors | TPR | FPR |
| --- | --- | --- | --- |
| FPCM | grad | 97.8333 | 7.5493 |
|  | mag | 98 | 8.6708 |
| Wavelet decomposition | grad | 96.6667 | 11.4282 |
|  | mag | 96.8333 | 12.4438 |
| Template matching | grad | 96.8333 | 12.3226 |
|  | mag | 96.3333 | 16.5424 |

The localization of the spikes detected by each method demonstrated spatial patterns corresponding to three modeled neuronal sources (see Fig. 12). The clusters obtained using FPCM detected spikes show somewhat better agreement with locations of the simulated sources of interictal activity than those found using the spikes detected with two other approaches.

**Figure 12:**
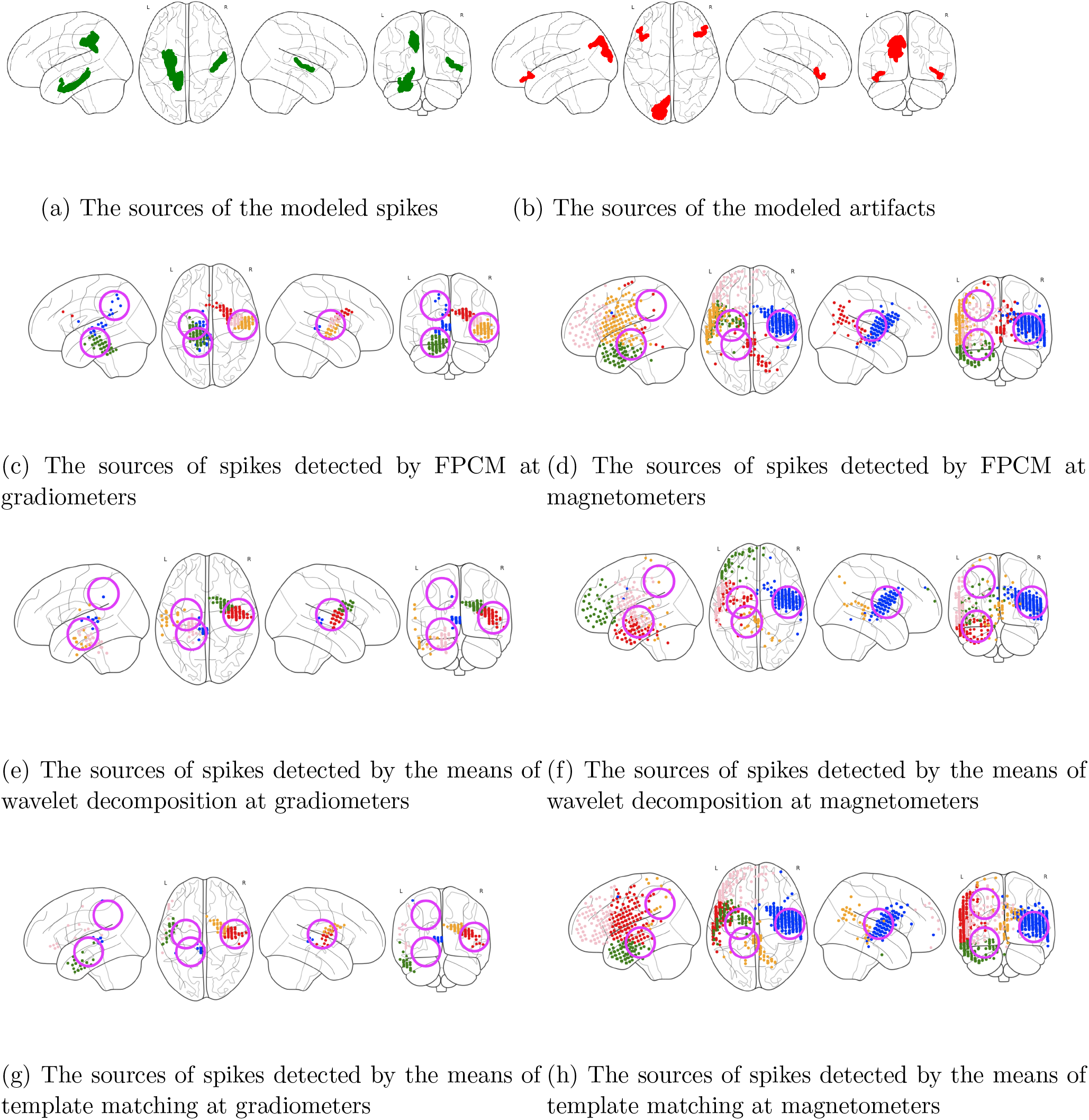
The comparison of spike localization performed by FPCM, wavelet decomposition or template matching at different types of sensors. The pink circles indicate the regions of the true spike generators. The colors of the sources indicate the cluster index obtained by K-means clustering of sources coordinates.

#### 3.1.2. ECoG (open dataset)

We applied FPCM, wavelet decomposition and template matching to ECoG simulations from (Quiroga et al., 2004). As evident from the Table 4, FPCM appears to be more stable to the noise level increase than the other two approaches. This behavior is in line with the observations based on our own simulated MEG data, see Fig. 10. The greater superiority of the FPCM solution in our data as compared to the behavior it exhibits with (Quiroga et al., 2004) dataset can be explained by the non-stationarity of noise in our simulations. Nevertheless in low SNR cases the FPCM appears to be superior to the other two approaches tested.

**Table 4:** The performance of FPCM, wavelet decomposition and template matching at different noise levels of simulated ECoG data from (Quiroga et al., 2004)

| Metrics | Noise level | FPCM | Wavelet decomp. | Template matching |
| --- | --- | --- | --- | --- |
| ROC AUC | 0.05 | 98.1797 | 99.9792 | 97.8194 |
|  | 0.1 | 98.0772 | 99.8491 | 97.4352 |
|  | 0.2 | 97.8708 | 98.6405 | 98.2474 |
|  | 0.3 | 97.9058 | 96.9925 | 90.518 |
|  | 0.4 | 97.8742 | 95.4728 | 95.8434 |
| PR AUC | 0.05 | 86.1303 | 99.8267 | 90.0194 |
|  | 0.1 | 85.8414 | 99.2655 | 88.9519 |
|  | 0.2 | 84.8966 | 94.398 | 91.0469 |
|  | 0.3 | 84.9678 | 87.5663 | 64.4028 |
|  | 0.4 | 84.6919 | 80.2929 | 75.2997 |

### 3.2. Real data (humans)

The analysis of the real data from three patients allowed to localize distinguishable areas, which are potentially epileptogenic (see Fig. 13, 14 and 15). For the patient A the dipole clusters were found bilaterally in the two cortical regions which matched manual analysis by an epileptologist, for the patient B – in the right temporal lobe, while for patient C – in the left temporal and parietal areas. For the patient B the resection was performed in the anterior part of the right temporal lobe, resulting in the seizure-free outcome several months later. The patient C underwent two resections. The first resection performed was not entirely successful as it included only the anterior portion of the left temporal lobe. Subsequent reevaluation of the patient detected spikes in the posterior part of the temporal lobe and the patient was sent to gamma-knife surgery to remove the posterior portion of the temporal lobe which resulted in the patient being seizure free on medications for several months.

**Figure 13:**
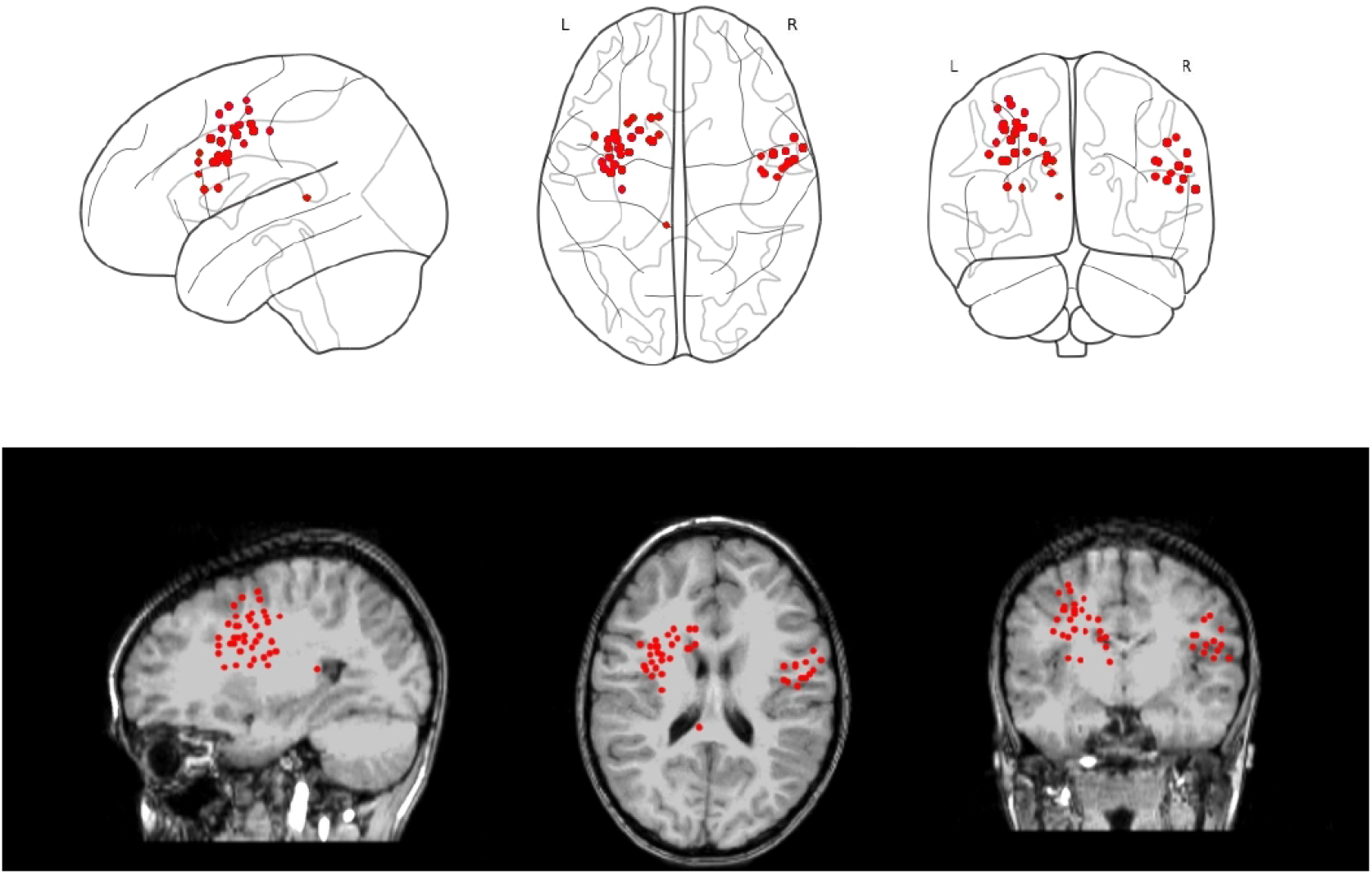
RAP-MUSIC localization of the FPCM detected interictal spikes in patient A displayed schematically and on the individual MRI.

**Figure 14:**
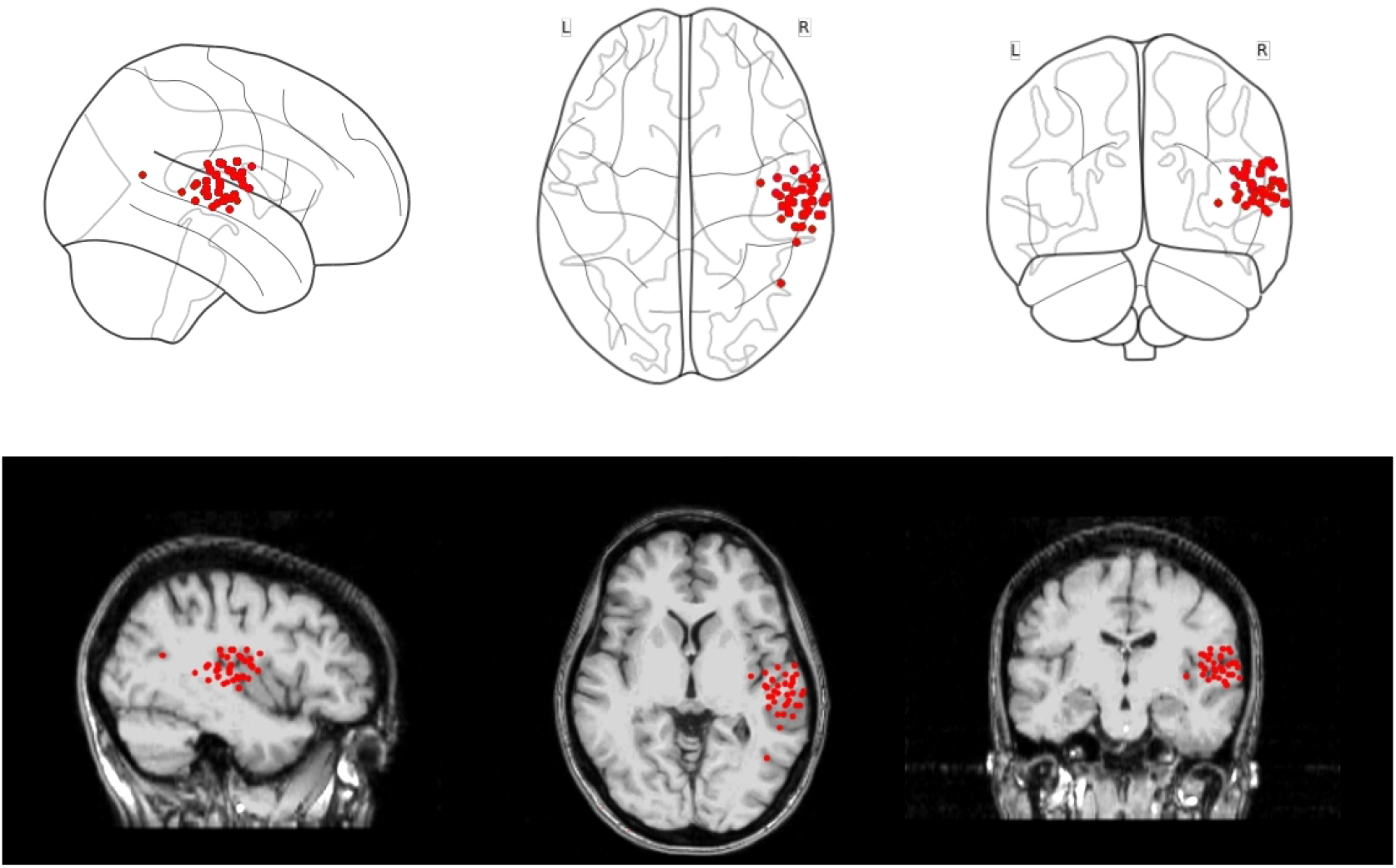
RAP-MUSIC localization of the FPCM detected interictal spikes in patient B displayed schematically and on the individual MRI.

**Figure 15:**
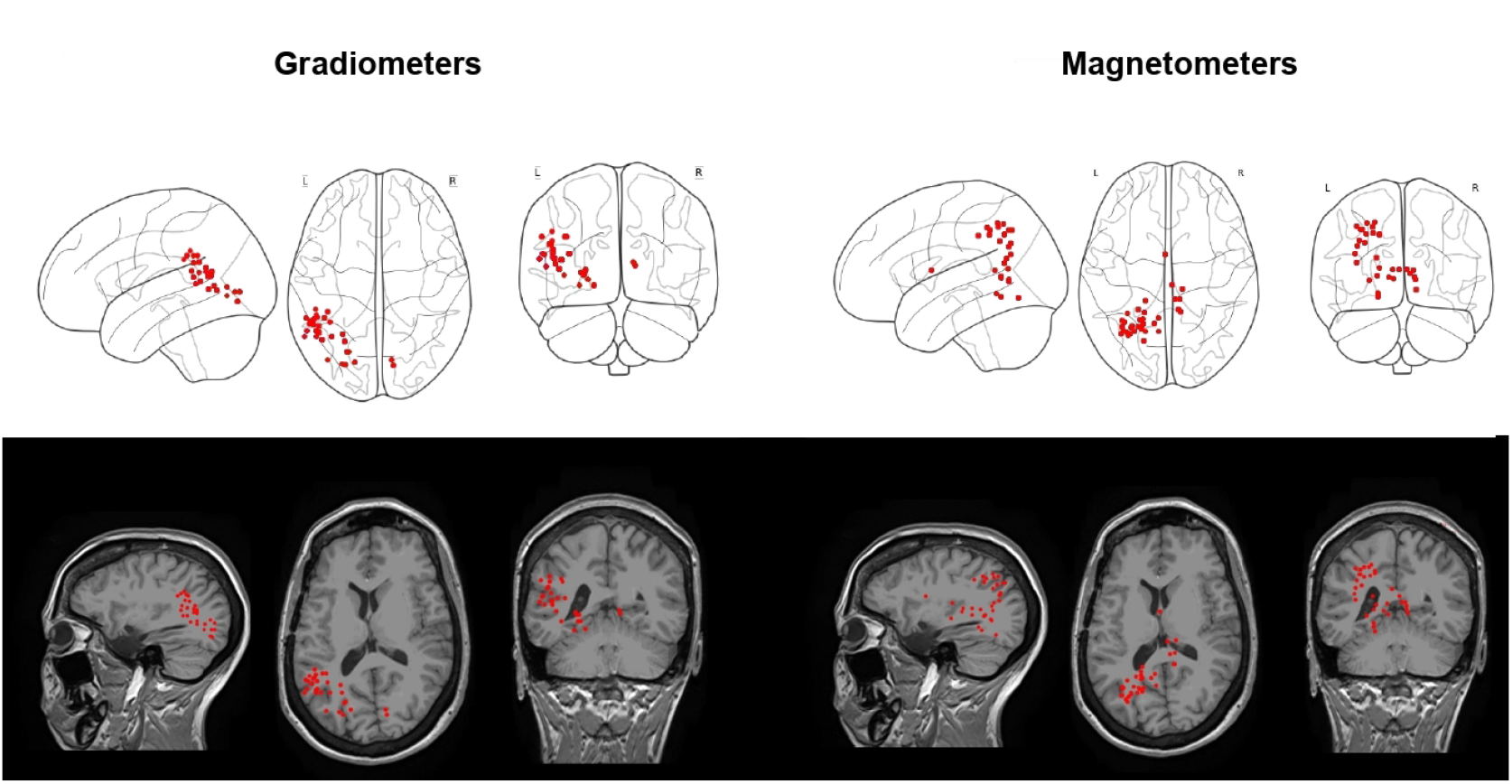
RAP-MUSIC localization of the FPCM detected interictal spikes in patient C displayed schematically and on the individual MRI.

The discrepancies in locations of the spikes detected from gradiometers and magnetometers in patient C reveal that gradiometers are less contaminated by the activity from deep sources and may produce the results with tighter clusters.

### 3.3. Noisy human EEG data

In order to demonstrate robustness of the FPCM technique against spontaneous high amplitude transients we applied FPCM and the other two standard approaches to the EEG data recorded in a patient at an ICU. This short dataset comprised 12 channels with total 11 manually detected spikes at 2 distinct latency values distributed over several channels (see Fig. 16).

**Figure 16:**
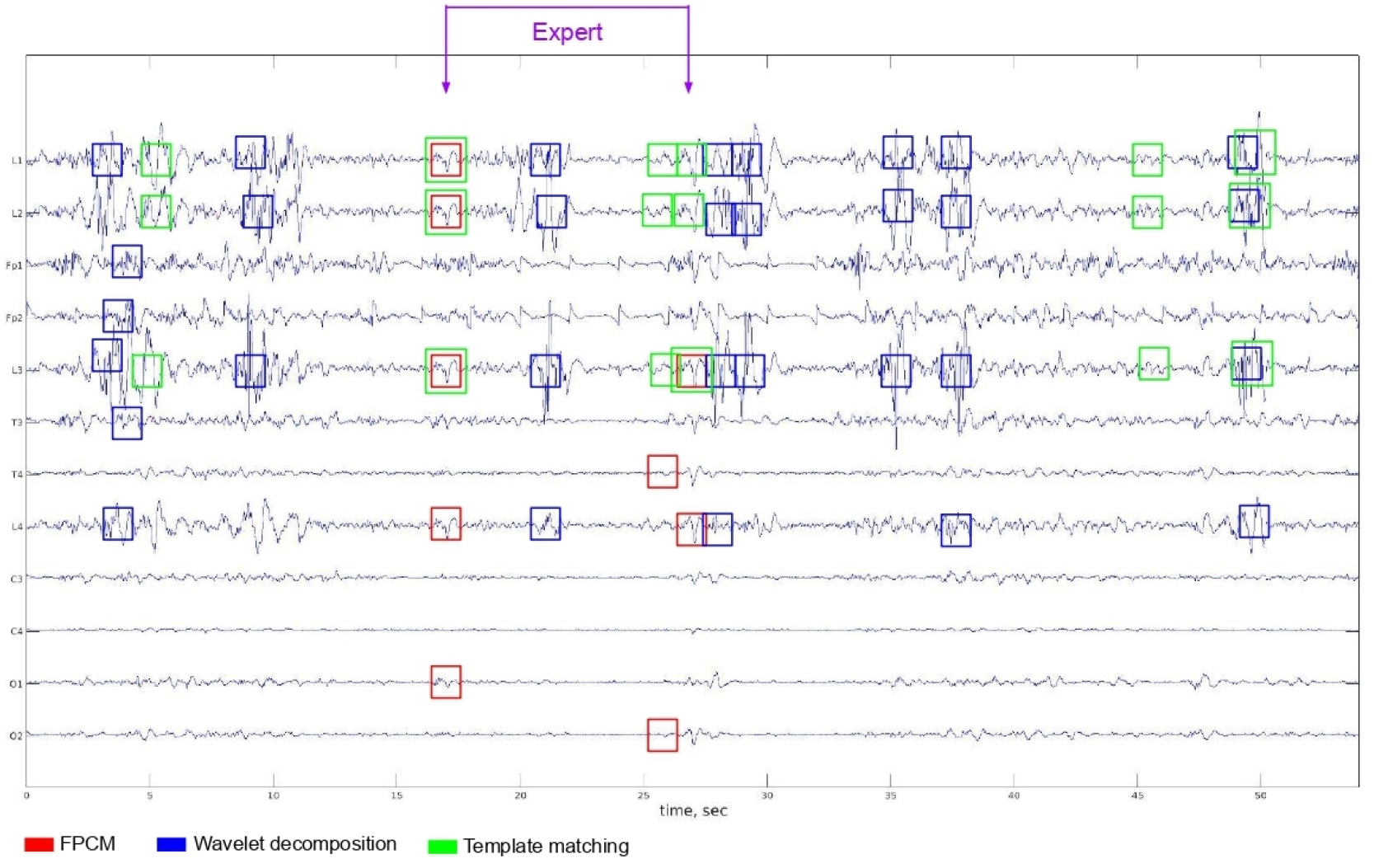
The comparison of patterns detected by the means of FPCM, wavelet decomposition and template matching. The EEG interictal peak-wave complexes are marked by the expert (purple arrows). The timeseries also exhibit large high frequency transients that obscure significantly less salient interictal events.

The proposed FPCM technique detected 7 true spikes and 2 false spikes which resulted into catching both true spike time moments and one false positive over 1 minute of data. Note that no threshold on absolute spike magnitude was used here. The demonstrated sensitivity is thus 77.78% if the task is to detect spikes in each channel independently and 100% if we aim at detecting the time moments when the spikes happen. Note that FPCM also detected one extra tiny event due to the fact that FPCM operates purely based on the shape parameter and not the absolute magnitude.

Wavelet-based approach appeared to be too sensitive to the sharp artifacts and failed to detect any of the true spikes (sensitivity was equal to 0 %). At the same time it marked 63 false patterns as spikes. The number of false positive latencies was equal to 21 within 1 minute.

For template matching we used several templates based on the spikes previously detected by FPCM. Although template matching succeeded to detect some of them (4 true positives), it missed 7 spikes and erroneously marked as spikes 41 segments. The algorithm covered both true positive latencies, but detected 13 false positives. The resulting sensitivity appeared to be equal to 36.36 %.

The purpose of the above presentation is to solely demonstrate the robustness of the proposed approach in the presence of high amplitude transients within frequency content similar to that of the target events. As we show this confirms the observations with simulated data and exhibits the similar of the three different methods with respect to the tolerance to high amplitude fast transients. The performance of all the wavelet and template based detectors could clearly be improved by preprocessing the data using spatial and temporal filtering techniques. At the same time the proposed approach will also benefit from such data conditioning.

### 3.4. Real data (rats)

To demonstrate the versatility of the proposed technique we have also applied it to a 20 s long segment of EEG data collected from rats who developed epileptic activity following the TBI. First the data were manually marked by an expert to comprise 39 interictal events. In this data the FPCM detected total of 61 spikes (see Fig. 17) missing only 3 out of 39 manually marked events. Overall the FPCM algorithm demonstrated 92.3% TPR and 59% FPR.

**Figure 17:**
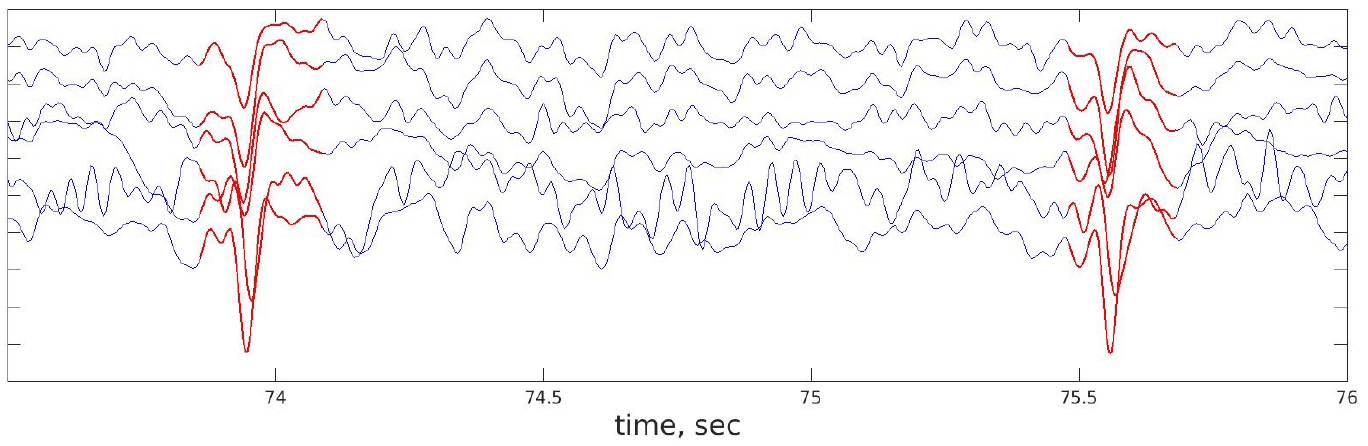
FPCM detected spikes. Here we show only a 3 seconds long segment of the time series. The intervals colored with red highlight the detected spikes.

Qualitative analysis of marked splines illustrates that false positive spikes can be divided into two groups: false positives with opposite polarity (apex points upwards) and negative polarity. These spikes despite the appropriate shape, are of low-amplitude and therefore an additional criteria based on the spike magnitude could further improve the situation. We have also applied the basic K-means plus-plus approach to spikes found by the FPCM where vector of coefficients **c** was used as a feature. As a result, each potential spike was assigned to three clusters based on its morphological properties. The mean for each cluster is represented in Fig. 18. As we can see the spikes in the three clusters differed primarily in their magnitude. The cluster with the largest magnitude (3rd cluster) appeared to include the largest number of spikes (29) detected by an expert. The second largest cluster (2nd cluster) contained 7 manually detected events and the least powerful cluster contained only 1 such spike. Based on this analysis, we can clearly observe the expert’s bias towards high amplitude spikes for greater confidence.

**Figure 18:**
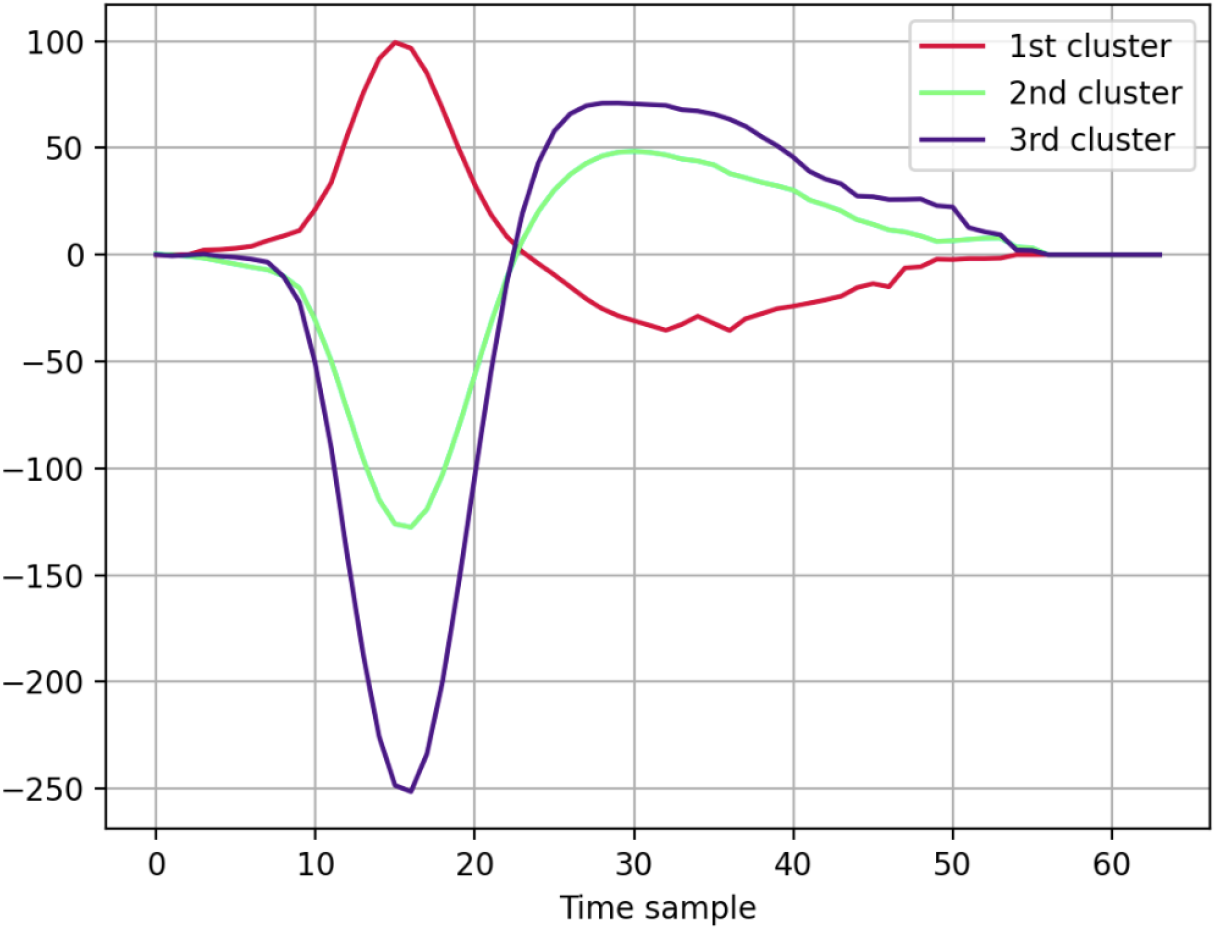
Means of the detected spike clusters. Spikes detected by FPCM were divided into three clusters using K-means plus-plus clustering procedure applied to the spline coefficient vectors. Then, the resulting waveforms were averaged within each cluster to obtain the depicted curves. The 1st cluster might represent the group of false positives with positive polarity, the 2nd cluster – false positives with negative polarity, 3rd cluster – group of actual spikes.

Removing the cluster with the smallest amplitude resulted into a slight decrease in TPR (90%) accompanied by a significant growth in FPR (71%). Thus, we can conclude that the FPCM approach is sufficiently versatile and can be successfully applied for detection of interictal discharges in animal data with a vast space for additional improvements and modifications based upon the basic idea presented here.

ROC analysis was consistent with the results previously reported with the use of simulated data: the largest ROC AUC corresponded to FPCM(ROC AUC = 97.475). The performance of wavelet decomposition appeared to be slightly worse (ROC AUC=91.228), and the worst performance in this task pertains to template matching (ROC AUC=77.4817) (see Fig. 19). It should be noted that FPCM and template matching reach high levels of TPR at low FPR (less than 5 × 10^-3^). Template matching reaches the same level as well, but at FPR=0.08.

**Figure 19:**
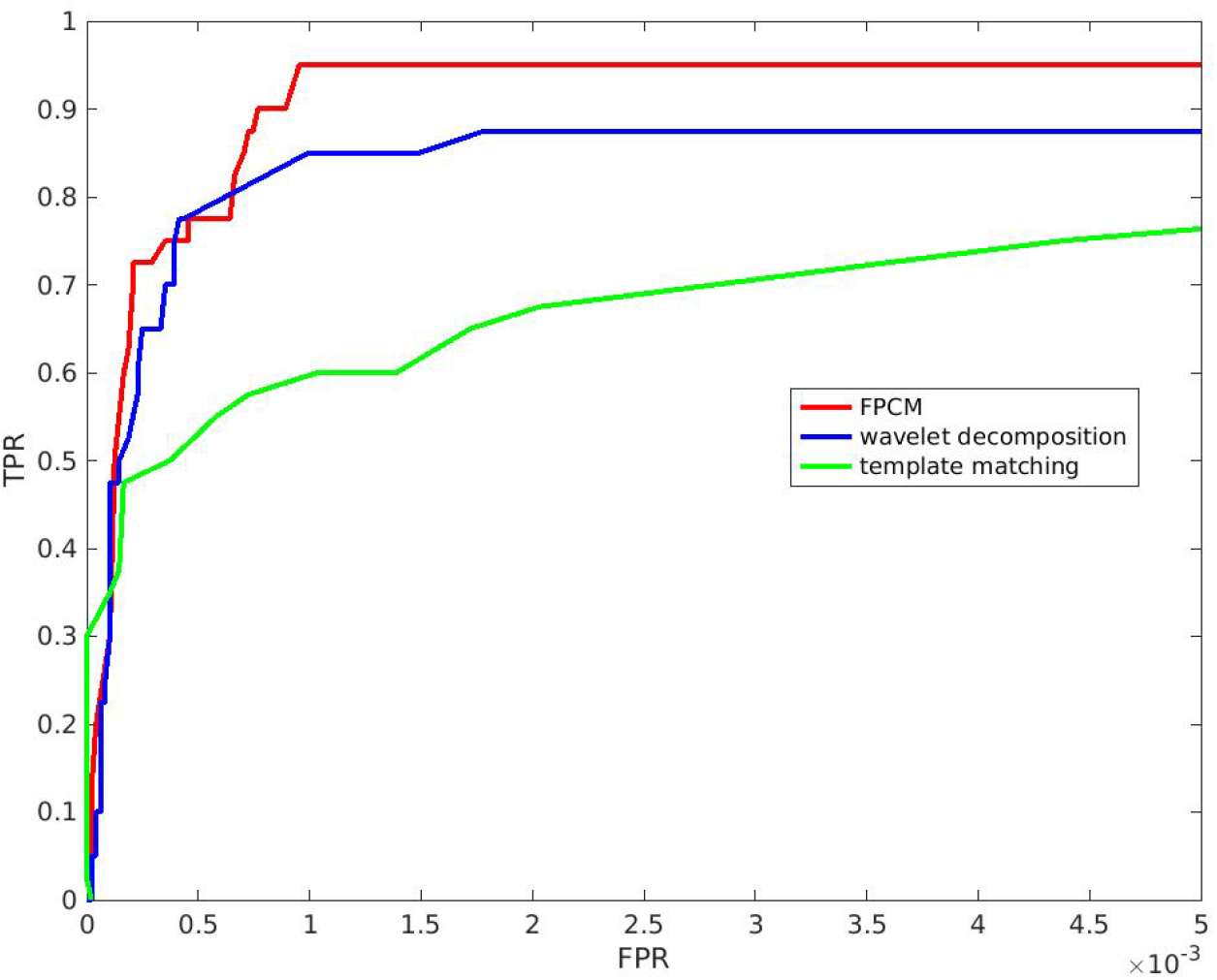
ROC curves comparing the performance of FPCM with that of the standard methods applied to the data collected from rats

## 4. Summary and conclusion

We have described a new mimetic approach for detection of interictal spikes in the electrophysiological data. This method generalizes the mixed spline machinery and builds the analytic morphological model of the target events. The resulting model comprises two linear and one parabolic segments whose exact shape depends on set of six mixed spline coefficients. As we show these coefficients can be effortlessly estimated for each time sample allowing for a quick and easily vectorized test on the morphology of the resulting mixed spline model applied at each data point. To do so, a verbal description of the expected target event morphology is turned into a set of logical predicates exploiting signs and ratios of the six spline coefficients. Fulfillment of these predicates in combination with a small percentage of the unexplained variance supports conclusions about the presence of the target event at a given latency. The approach is applicable to detection of not only interictal spikes that can be well described by a pair of segments. The number of segments and the corresponding polynomials can be chosen to describe the target shape and the model can be recreated according to the principles outlined earlier.

Our mimetic technique relies on the computationally light convolution operation which allows for rapid scanning of the large amount of data or for a real-time application during patients monitoring session. Also, the FPCM approach can be successfully applied for detection of pathological interictal activity in animal recordings. The basic methodology described here can easily adopt other more complex shapes and perform rapid scanning for the presence of, for example, K-complexes, ECG or alternative specific shapes in the electrophysiological data other than that collected from epileptic brains, including neural spikes. Speaking of detection of spikes produced by individual neurons, which can be characterized by overlapping and other specific properties, there is a variety of neural spikes detection methods such as teager energy operation (Kim and Kim, 2000), approaches based on stochastic resonance (Güngör and Töreyin, 2020), sparse signal decomposition (Ekanadham et al., 2014). The most recent techniques include normalized template matching (Laboy-Juárez et al., 2019) and deep-learning based methods (Saif-ur Rehman et al., 2019). At present FPCM can not be claimed to be superior to the well-established techniques of neural spikes detection observed in the intracellular recordings. However, there are crucial differences between this task and the task of detection of interictal transients. Individual neurons’ spike shape variability is less than that observed in the interictal discharges produced by superposition of activity of a large number of neurons in a pathological brain. Also, historically, neuronal spikes were detected using automatic methods while interictal spikes traditionally have been identified visually based on a set of guidelines Adjouadi et al. (2004) that get refined every now and then. This makes it important to be able to perform automatic detection using a biomimetic approach so that the guidelines adopted by the human experts can be easily passed to the algorithm. Also, such biomimetic algorithms are likely to be more easily translated into a clinical practice since they naturally implement well established spike identification criteria accepted in the clinical community. We expect that the flexibility of FPCM’s parameters and its low computational demands warrants exploring its applicability the tasks other than interictal spike detection explored in this paper.

Throughout the paper we compared our approach to two basic methods. The first reference technique relies on using wavelet coefficients as features for detection of interictal events. While promising this method appeared to be sensitive to large amplitude transients and therefore unlike our approach requires additional heuristics including data preprocessing to operate properly. Another reference methodology is based on template matching which is basically match filtering known to be optimal in case of white additive noise. Frequency domain whitening could potentially improve the performance here. However, given that spike spectra overlaps with the spectral properties of the ongoing brain activity such whitening would inevitably lead to the decreased SNR of individual spikes. Also, the inherent non-stationarity of EEG and MEG data compromises the performance of such a whitening step. These two arguments lead us to not using any whitening procedure in our implementation of the template matching technique. It is also noteworthy that FPCM relies on the convolution operation and in fact performs parametric template matching. However, our method is equipped with an additional flexibility that lies in the mimetic decision making based on a set of logical predicates. This combination provides the extra robustness that FPCM enjoys when compared to the classical template based matching. These properties of the proposed technique resulted in FPCM being immune to the presence of the high amplitude artifacts, outperforming wavelet decomposition and template matching.

The FPCM technique can be applied to either raw timeseries of independent components to improve the results reported in Ossadtchi et al. (2004); Kobayashi et al. (2002a). At the same time, the use of spatial decomposition methods followed by the components selection process may introduce additional errors in cases extra components are kept or some pivotal components are omitted. Therefore, primarily we consider the FPCM as an alternative approach to be applied to raw data or the data where spatial decomposition is used for denoising and not for the selection of a small number of target components containing interictal spikes. Robustness of the FPCM with respect to high amplitude transients makes it a perfect candidate for such a use.

The described technique can be further extended and improved. In order to compute the mixed spline coefficients we solve an over-determined system of linear equations in the least squares sense. This means that we implicitly assume normality of the residuals and our methodology becomes suboptimal when this assumption is violated. In reality the ongoing spontaneous EEG activity is often not normally distributed and may be described by a heavy-tailed distribution. Therefore, our approach would benefit from taking into account the potential non-Gaussianity of the background data. This can be accomplished for example in a fashion similar to that described in Jas et al. (2017).

Another limitation of the proposed methodology is the requirement for the elementary segments to be necessarily described by a polynomial function. Although theoretically we can approximate any function in the vicinity of a specific point using Taylor series expansion, for complex shapes this would require many terms and will increase the number of convolutions to be performed in order to estimate mixed spline coefficients. Also, forming logical predicates for the coefficients of polynomials with a large number of terms becomes less intuitive. This means that FPCM suites best for detection of relatively simple shapes where each elementary segment is described by either linear, quadratic or cubic polynomial.

Despite the listed limitations the proposed FPCM technique represents a viable method to automate the visual search procedure traditionally used in exploration of interictal data. Unlike several other approaches like Ossadtchi et al. (2004) FPCM does not rely on a spatial decomposition and thus reduces the risks of missing the entire cluster corresponding to a potentially epileptogenic zone. Its low computational requirements and tolerance to artifacts make it an ideal candidate for a robust against outliers real-time detection of events with specific morphology. Finally, FPCM’s versatility allows for applications beyond monitoring of epilepsy brains in humans and animals, the avenues we hope to explore in the future.

## 5. Acknowledgement

This work is supported by the Center for Bioelectric Interfaces NRU HSE, RF Government grant, AG. No. 075-15-2021-624. We thank Dr. Tommaso Fedele for fruitful discussions and references to the relevant literature and Dr. Mikhail Lebedev for funding acquisition.

## 6. Appendix

**Table 5:** AUC values comparing the performance of different methods at different SNRs

| Sensors | Artifacts' amplitude | FPCM | Wavelet decomp. | Template matching |
| --- | --- | --- | --- | --- |
| grad | 0 | 98.2494 | 98.6607 | 98.4946 |
|  | 35 | 98.2469 | 98.4915 | 98.3207 |
|  | 70 | 98.2441 | 97.9853 | 97.7368 |
|  | 105 | 98.1579 | 97.1020 | 96.9872 |
|  | 140 | 98.0803 | 96.1413 | 96.5435 |
| mag | 0 | 98.4339 | 99.1337 | 98.4154 |
|  | 35 | 98.4117 | 98.9633 | 98.4855 |
|  | 70 | 98.2870 | 98.3338 | 97.3692 |
|  | 105 | 98.0803 | 97.3614 | 94.9678 |
|  | 140 | 97.9310 | 96.3959 | 92.9672 |

## References

Abd El-Samie, F. E., Alotaiby, T. N., Khalid, M. I., Alshebeili, S. A., and Aldosari, S. A. (2018). A review of eeg and meg epileptic spike detection algorithms. IEEE Access, 6:60673–60688.

Adjouadi, M., Sanchez, D., Cabrerizo, M., Ayala, M., Jayakar, P., Yaylali, I., and Barreto, A. (2004). Interictal spike detection using the walsh transform. IEEE Transactions on Biomedical Engineering, 51(5):868–872.

Calvagno, G., Ermani, M., Rinaldo, R., and Sartoretto, F. (2000). A multiresolution approach to spike detection in eeg. In 2000 IEEE International Conference on Acoustics, Speech, and Signal Processing. Proceedings (Cat. No. 00CH37100), volume 6, pages 3582–3585. IEEE.

De Oliveira, P. G., Queiroz, C., and Da Silva, F. L. (1983). Spike detection based on a pattern recognition approach using a microcomputer. Electroencephalography and clinical neurophysiology, 56(1):97–103.

Dingle, A. A., Jones, R. D., Carroll, G. J., and Fright, W. R. (1993). A multistage system to detect epileptiform activity in the eeg. IEEE Transactions on Biomedical Engineering, 40(12):1260–1268.

Ekanadham, C., Tranchina, D., and Simoncelli, E. P. (2014). A unified framework and method for automatic neural spike identification. Journal of neuroscience methods, 222:47–55.

El-Gohary, M., McNames, J., and Elsas, S. (2008). User-guided interictal spike detection. In 2008 30th Annual International Conference of the IEEE Engineering in Medicine and Biology Society, pages 821–824. IEEE.

Faure, C. (1985). Attributed strings for recognition of epileptic transients in eeg. International journal of bio-medical computing, 16(3-4):217–229.

Glover, J. R., Ktonas, P. Y., Raghavan, N., Urunuela, J. M., Velamuri, S. S., and Reilly, E. L. (1986). A multichannel signal processor for the detection of epileptogenic sharp transients in the eeg. IEEE transactions on biomedical engineering, (12):1121–1128.

Glover, J. R., Raghaven, N., Ktonas, P. Y., and Frost, J. D. (1989). Context-based automated detection of epileptogenic sharp transients in the eeg: elimination of false positives. IEEE Transactions on Biomedical Engineering, 36(5):519–527.

Gotman, J. and Gloor, P. (1976). Automatic recognition and quantification of interictal epileptic activity in the human scalp eeg. Electroencephalography and clinical neurophysiology, 41(5):513–529.

Gotman, J. and Wang, L. (1991). State-dependent spike detection: concepts and preliminary results. Electroencephalography and clinical Neurophysiology, 79(1):11–19.

Gotman, J. and Wang, L.-Y. (1992). State dependent spike detection: validation. Electroencephalography and clinical neurophysiology, 83(1):12–18.

Gramfort, A., Luessi, M., Larson, E., Engemann, D. A., Strohmeier, D., Brodbeck, C., Parkkonen, L., and Hämäläinen, M. S. (2014). Mne software for processing meg and eeg data. Neuroimage, 86:446–460.

Güngör, C. B. and Töreyin, H. (2020). Facilitating stochastic resonance as a pre-emphasis method for neural spike detection. Journal of Neural Engineering, 17(4):046047.

Halford, J. J., Schalkoff, R. J., Zhou, J., Benbadis, S. R., Tatum, W. O., Turner, R. P., Sinha, S. R., Fountain, N. B., Arain, A., Pritchard, P. B., et al. (2013). Standardized database development for eeg epileptiform transient detection: Eegnet scoring system and machine learning analysis. Journal of neuroscience methods, 212(2):308–316.

Hostetler, W. E., Doller, H. J., and Homan, R. W. (1992). Assessment of a computer program to detect epileptiform spikes. Electroencephalography and clinical neurophysiology, 83(1):1–11.

Indiradevi, K., Elias, E., Sathidevi, P., Nayak, S. D., and Radhakrishnan, K. (2008). A multi-level wavelet approach for automatic detection of epileptic spikes in the electroencephalogram. Computers in biology and medicine, 38(7):805–816.

Jas, M., La Tour, T. D., Şimşekli, U., and Gramfort, A. (2017). Learning the morphology of brain signals using alpha-stable convolutional sparse coding. arXiv preprint arXiv:1705.08006.

Ji, Z., Sugi, T., Goto, S., Wang, X., Ikeda, A., Nagamine, T., Shibasaki, H., and Nakamura, M. (2011). An automatic spike detection system based on elimination of false positives using the large-area context in the scalp eeg. IEEE transactions on biomedical engineering, 58(9):2478–2488.

Keshri, A. K., Sinha, R. K., Singh, A., and Das, B. N. (2011). Dfaspike: A new computational proposition for efficient recognition of epileptic spike in eeg. Computers in biology and medicine, 41(7):559–564.

Kim, K. H. and Kim, S. J. (2000). Neural spike sorting under nearly 0-db signal-to-noise ratio using nonlinear energy operator and artificial neural-network classifier. IEEE Transactions on Biomedical Engineering, 47(10):1406–1411.

Kim, S. and McNames, J. (2007). Automatic spike detection based on adaptive template matching for extracellular neural recordings. Journal of neuroscience methods, 165(2):165–174.

Kobayashi, K., Akiyama, T., Nakahori, T., Yoshinaga, H., and Gotman, J. (2002a). Systematic source estimation of spikes by a combination of independent component analysis and rap-music: I: Principles and simulation study. Clinical Neurophysiology, 113(5):713–724.

Kobayashi, K., Akiyama, T., Nakahori, T., Yoshinaga, H., and Gotman, J. (2002b). Systematic source estimation of spikes by a combination of independent component analysis and rap-music: Ii: preliminary clinical application. Clinical neurophysiology, 113(5):725–734.

Komoltsev, I. G., Sinkin, M. V., Volkova, A. A., Smirnova, E. A., Novikova, M. R., Kordonskaya, O. O., Talypov, A. E., Guekht, A. B., Krylov, V. V., and Gulyaeva, N. V. (2020). A translational study on acute traumatic brain injury: high incidence of epileptiform activity on human and rat electrocorticograms and histological correlates in rats. Brain sciences, 10(9):570.

Kural, M. A., Duez, L., Hansen, V. S., Larsson, P. G., Rampp, S., Schulz, R., Tankisi, H., Wennberg, R., Bibby, B. M., Scherg, M., et al. (2020). Criteria for defining interictal epileptiform discharges in eeg: A clinical validation study. Neurology, 94(20):e2139–e2147.

Laboy-Juárez, K. J., Ahn, S., and Feldman, D. E. (2019). A normalized template matching method for improving spike detection in extracellular voltage recordings. Scientific reports, 9(1):1–12.

Latka, M., Was, Z., Kozik, A., and West, B. J. (2003). Wavelet analysis of epileptic spikes. Physical Review E, 67(5):052902.

Liu, Y.-C., Lin, C.-C. K., Tsai, J.-J., and Sun, Y.-N. (2013). Model-based spike detection of epileptic eeg data. Sensors, 13(9):12536–12547.

Lodder, S. S., Askamp, J., and van Putten, M. J. (2013). Inter-ictal spike detection using a database of smart templates. Clinical neurophysiology, 124(12):2328–2335.

Maragos, P. and Schafer, R. (1987). Morphological filters–part i: Their set-theoretic analysis and relations to linear shift-invariant filters. IEEE Transactions on Acoustics, Speech, and Signal Processing, 35(8):1153–1169.

Mosher, J. C. and Leahy, R. M. (1999). Source localization using recursively applied and projected (rap) music. IEEE Transactions on signal processing, 47(2):332–340.

Nishida, S., Nakamura, M., Ikeda, A., and Shibasaki, H. (1999). Signal separation of background eeg and spike by using morphological filter. Medical engineering & physics, 21(9):601–608.

Nonclercq, A., Foulon, M., Verheulpen, D., De Cock, C., Buzatu, M., Mathys, P., and Van Bogaert, P. (2009). Spike detection algorithm automatically adapted to individual patients applied to spike and wave percentage quantification. Neurophysiologie Clinique/Clinical Neurophysiology, 39(2):123–131.

Nonclercq, A., Foulon, M., Verheulpen, D., De Cock, C., Buzatu, M., Mathys, P., and Van Bogaert, P. (2012). Cluster-based spike detection algorithm adapts to interpatient and intrapatient variation in spike morphology. Journal of neuroscience methods, 210(2):259–265.

Ossadtchi, A., Baillet, S., Mosher, J., Thyerlei, D., Sutherling, W., and Leahy, R. (2004). Automated interictal spike detection and source localization in magnetoencephalography using independent components analysis and spatio-temporal clustering. Clinical Neurophysiology, 115(3):508–522.

Pon, L.-S., Sun, M., and Sclabassi, R. J. (2002). The bi-directional spike detection in eeg using mathematical morphology and wavelet transform. In 6th International Conference on Signal Processing, 2002., volume 2, pages 1512–1515. IEEE.

Quiroga, R. Q., Nadasdy, Z., and Ben-Shaul, Y. (2004). Unsupervised spike detection and sorting with wavelets and superparamagnetic clustering. Neural computation, 16(8):1661–1687.

Saif-ur Rehman, M., Lienkämper, R., Parpaley, Y., Wellmer, J., Liu, C., Lee, B., Kellis, S., Andersen, R., Iossifidis, I., Glasmachers, T., et al. (2019). Spikedeeptector: a deep-learning based method for detection of neural spiking activity. Journal of neural engineering, 16(5):056003.

Sankar, R. and Natour, J. (1992). Automatic computer analysis of transients in eeg. Computers in biology and medicine, 22(6):407–422.

Sartoretto, F. and Ermani, M. (1999). Automatic detection of epileptiform activity by single-level wavelet analysis. Clinical Neurophysiology, 110(2):239–249.

Scheuer, M. L., Bagic, A., and Wilson, S. B. (2017). Spike detection: Inter-reader agreement and a statistical turing test on a large data set. Clinical Neurophysiology, 128(1):243–250.

Senhadji, L., Dillenseger, J.-L., Wendling, F., Rocha, C., and Kinie, A. (1995). Wavelet analysis of eeg for three-dimensional mapping of epileptic events. Annals of biomedical engineering, 23(5):543–552.

Staley, K. J. and Dudek, F. E. (2006). Interictal spikes and epileptogenesis. Epilepsy Currents, 6(6):199–202.

Subasi, A. (2006). Automatic detection of epileptic seizure using dynamic fuzzy neural networks. Expert Systems with Applications, 31(2):320–328.

Webber, W. R. S., Litt, B., Lesser, R. P., Fisher, R., and Bankman, I. (1993). Automatic eeg spike detection: what should the computer imitate? Electroencephalography and clinical neurophysiology, 87(6):364–373.

Wilson, S. B. and Emerson, R. (2002). Spike detection: a review and comparison of algorithms. Clinical Neurophysiology, 113(12):1873–1881.

Xu, G., Wang, J., Zhang, Q., Zhang, S., and Zhu, J. (2007). A spike detection method in eeg based on improved morphological filter. Computers in biology and medicine, 37(11):1647–1652.

